# Glycocalyx crowding with synthetic mucin mimetics strengthens interactions between soluble and virus-associated lectins and cell surface glycan receptors

**DOI:** 10.1101/2021.05.07.443169

**Authors:** Daniel J. Honigfort, Meghan O. Altman, Pascal Gagneux, Kamil Godula

## Abstract

Membrane-associated mucins protect epithelial cell surfaces against pathogenic threats by serving as non-productive decoys that capture infectious agents and clear them from the cell surface and by erecting a physical barrier that restricts their access to target receptors on host cells. However, the mechanisms through which mucins function are still poorly defined due to a limited repertoire of tools available for tailoring their structure and composition in living cells with molecular precision. Using synthetic glycopolymer mimetics of mucins, we modeled the mucosal glycocalyx on red blood cells (RBC) and evaluated its influence on lectin (SNA) and virus (H1N1) adhesion to endogenous sialic acid receptors. The glycocalyx inhibited the rate of SNA and H1N1 adhesion in a size- and density-dependent manner, consistent with current view of the mucins as providing a protective shield against pathogens. Counterintuitively, increasing density of the mucin mimetics enhanced the retention of bound lectins and viruses. Careful characterization of SNA behavior at the RBC surface using a range of biophysical and imaging techniques revealed lectin-induced crowding and reorganization of the glycocalyx with concomitant enhancement in lectin clustering, presumably through the formation of a more extensive glycan receptor patch at the cell surface. Our findings indicate that glycan-targeting pathogens may exploit the biophysical and biomechanical properties of mucins to overcome the mucosal glycocalyx barrier.

**Significance:** Like other animal hosts, humans are constantly challenged by pathogens. This has led to an evolution of physical barriers coating all mucosal tissues, which are most vulnerable to infection. An important part of this defense is a dense brush of large proteins, called mucins, which are heavily decorated with sugars and keep pathogens at bay. Deciphering how pathogens overcome the mucin barrier is necessary to understand early stages of infection and to develop more effective treatments. By artificially installing the mucin-like shield on the surfaces of cells using synthetic sugar-bearing polymers, we have discovered a new physical mechanism by which proteins and viruses can exploit this barrier to more strongly adhere to their targets.

## Introduction

Many viral and bacterial pathogens that circulate in the human population, and continuously pose a risk of disease outbreaks and pandemics, utilize glycans on host cells to facilitate their adhesion and initiate infection.^1^ Sialic acids (Sias) represent one such cell surface glycan receptor frequently targeted by pathogens, such as the Influenza A virus (IAV) among others (**Fig 1a**). The Sia monosaccharide is typically added as a terminal modification on more complex glycans linked to membrane-associated proteins or glycolipids and as a result is prominently displayed at high concentrations on mammalian cells.^2^ Pathogen-associated proteins that recognize Sias, such as hemagglutinins (HA) in the case of IAVs, often share a low affinity binding site for the individual receptor structures but achieve high avidity binding to Sias displayed on cells through oligomerization and multivalency.^3,4,5^

**Figure 1.**
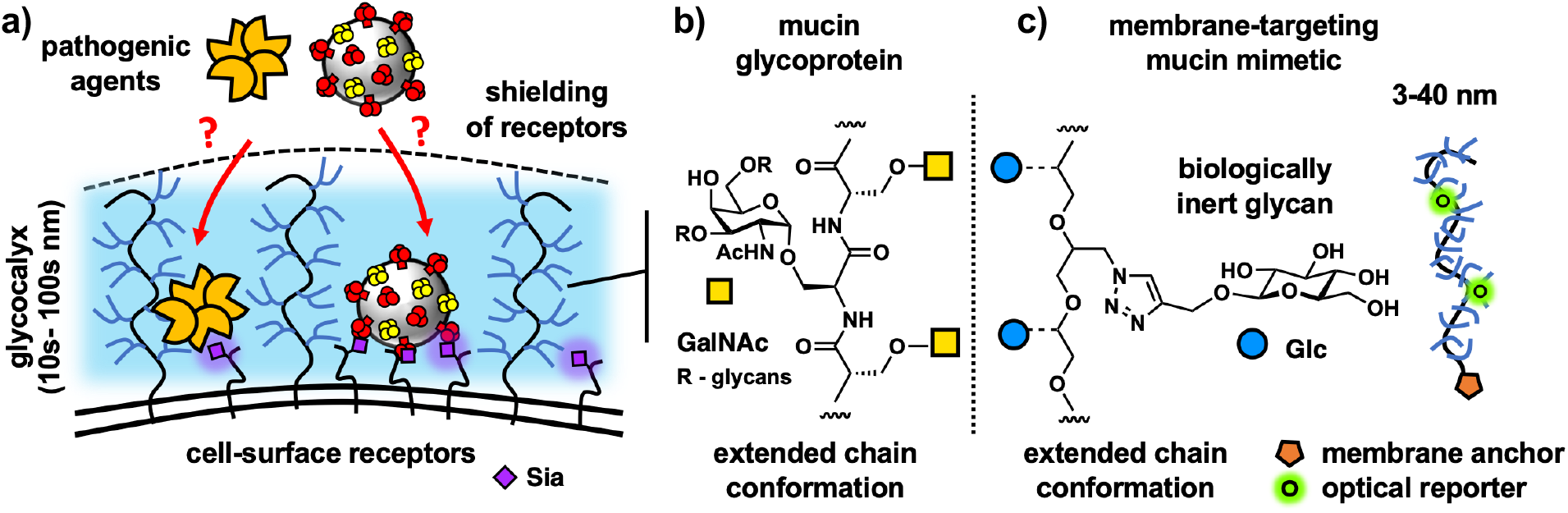
The mucosal glycocalyx provides a physical barrier against pathogen association with host cells. a) A brush of membrane-associated mucin glycoproteins restricts the ingress of pathogens toward cell surface glycan receptors, such as sialic acids (Sias). b) The extended conformation of mucins and their physical properties arise from dense glycosylation of their core peptide backbone. c) Cell surface targeting glycopolymers based on a linear poly(ethylene oxide) (PEO) scaffold, which mimic the architecture and physical properties of mucins, are used to investigate mechanisms through which pathogens can breach the protective mucosal glycocalyx barrier.

In response to the constant pathogen challenge, the cells of mucosal epithelial tissues, which are most susceptible to infection, express soluble and membrane bound glycoproteins called mucins to limit pathogen adhesion and spread (**Fig 1a**).^6,7^ Comprised of long polypeptide chains densely decorated with glycans (**Fig 1b**), membrane-associated mucins provide a dual protective function against infection by forming a dense extended biopolymer brush, restricting access to the cell surface and by presenting decoy receptors to capture and clear the pathogen.^8,9^ Both the biophysical barrier^10^ and the receptor decoy^11^ functions of mucins have been shown to limit bacterial and viral infection, including by IAVs. While the interactions between pathogens and mucins based on glycan recognition have received substantial attention, the biophysical mechanisms allowing mucins to limit infection are yet to be examined experimentally in a systematic way. This is primarily due to the lack of tools for manipulating the architecture and glycosylation status of mucins with molecular precision in living cells. Membrane engineering with synthetic glycopolymers, which mimic the architecture and properties of mucins and can be chemically defined at the molecular level,^12,13^ has emerged as a convenient approach to study phenomena associated with the mucin glycocalyx, including its interactions with sialic acid-binding proteins^14^ and the effects of its biophysical properties on cellular interactions and functions. ^15,16^ We have recently observed that introducing mucin mimetics to the surfaces of cells resulted in increased glycocalyx crowding, which limited lectin binding to membrane-associated glycoconjugates.^17^ Bertozzi and co-workers further showed that mucin mimetics based on synthetic glycopeptides imbedded in supported lipid bilayers could shield sialylated glycosphingolipids from H1N1.^18^ Studies with synthetic mucin analogs can reveal mechanistic insights into how mucins may limit pathogen adhesion through biophysical means.

To counteract the protective mucin shield, many pathogens express enzymes that degrade mucins^19,20^ or change their glycosylation status.^21^ While the complete removal of mucins by microbial proteases disrupts the integrity of the glycocalyx and exposes the cell surface to pathogen attack, the removal of only their receptor decoys with glycosidases, such as the IAV neuraminidase (NA), leaves the physical mucin barrier intact. The mechanism through which pathogens that do not break down the mucin shield overcome this physical barrier are yet to be fully investigated.

In this study, we have deployed synthetic glycopolymers with tunable lengths (**Fig 1c**) to model membrane-associate mucin displays on the surfaces of avian (turkey) red blood cells (RBCs) and examined how the mucin-mimetic glycocalyx size and density influenced the kinetics and thermodynamics of lectin (SNA) and viral (H1N1) binding to endogenous Sia receptors. We have observed polymer length-dependent inhibition of SNA and H1N1 adhesion to their cell surface receptors in the presence of the membrane-tethered mucin mimetic brush, providing further support for the mucin shield model. We have also identified a biophysical mechanism by which lectins and viruses exploit the crowded glycocalyx to strengthen their association with the cell.

## Results

### Glycocalyx engineering with mucin mimetics alters RBC morphology and properties

To model key physical features of the epithelial glycocalyx; i.e., its dense, extended brush-like organization of mucin glycoproteins, we have generated synthetic mucin-mimetic glycopolymers (**GPs**) with defined length and glycosylation (**Fig 1c**). The mucin-mimetic glycopolymers (**Fig 2a**) were assembled on linear hydrophilic poly(ethylene oxide) (PEO) scaffolds of increasing length (DP = 30, 140, 440) and with narrow dispersity (Đ < 1.3). The precursor backbones were generated via controlled anionic ring-opening polymerization of epichlorohydrin^22^ followed by functionalization with azide side-chains for the attachment of propargyl glycosides via copper-catalyzed azide-alkyne cycloaddition (CuAAc)^23^ (**Scheme S1**). The efficiency of this “click” process resulted in a high frequency of glycosylated side-chains in close proximity to the PEO backbone, forcing the glycopolymers into extended conformations characteristic of mucins.^17,24^ Seeking to isolate the biophysical properties of mucins from their glycan interactions, we modified our glycopolymers with glucose, a monosaccharide typically not found in mucin glycans (**Fig 1**). This produced mucin mimetic “spectators” lacking Sia receptors for IAVs but able to tune the size and density of the glycocalyx. The mucin mimetics were also endowed with a hydrophobic 4,5-*seco*-cholesten-5-one end-group for anchoring of the glycopolymers in cell membranes and a small fraction (< 1%) of their sidechains were labeled with AlexaFluor 488 (AF488) or Cyanine 3 (Cy3) dyes for detection via fluorescence (**Fig 1**). To examine the impact of glycocalyx size on molecular recognition events at the cell surface, we generated short (**GP-S**, DP = 30), medium (**GP-M**, DP=140), and long (**GP-L**, DP=440) mucin mimetics (**Fig 2A and 2B)**.^17^

**Figure 2.**
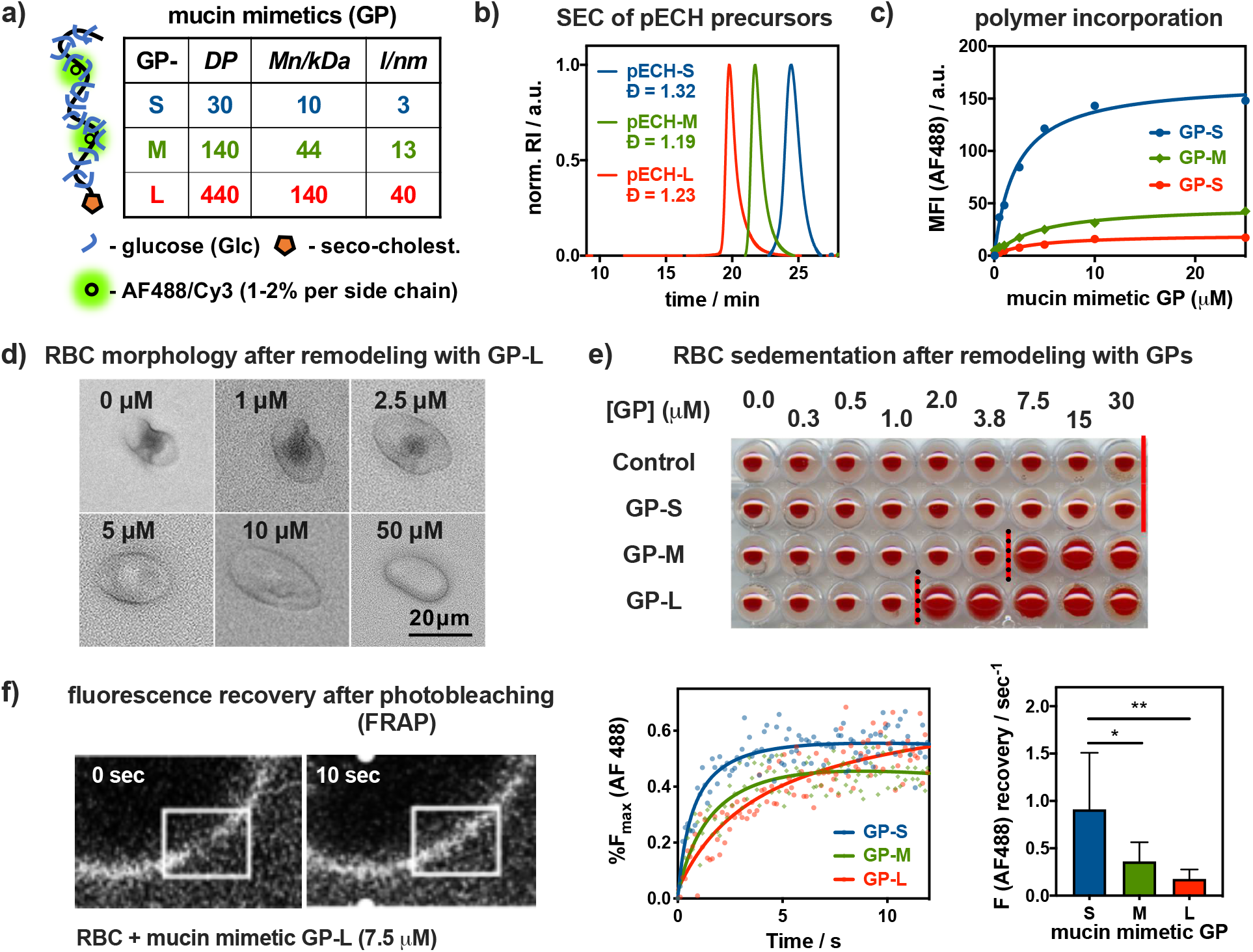
Construction and characterization of a mucosal glycocalyx model. **a)** Short (**S**), medium (**M**), and long (**L**) mucin mimetic glycopolymers **GP** ranging in size from ~ 3 to 40 nm were generated from poly(epichlorohydrin) (pECH) precursors (DP = degree of polymerization, *Mn* = number average molecular weight, *l* = estimated end to end length). **b)** Size exclusion chromatography (SEC) of pECH precursors indicates narrow molecular weight distributions of the polymer scaffolds. The polymers were modified with biologically inert glucose sidechains, fluorescent probes (AF488 or Cy3) for visualization, and chain end-terminated with hydrophobic cholestenone membrane anchor. **c)** Incorporation of AF488-labeled mucin mimetics **GPs** into RBC membranes was analyzed by flow cytometry. d) Optical microscopy and light scattering reveal increased cell rounding with increasing glycocalyx density (shown for **GP-L**). **e)** Sedimentation of RBCs remodeled with increasing concentration of all three glycopolymers (c_**GP**_ = 0.25 to 30 μM) **f)** Fluorescence recovery after photobleaching (FRAP) analysis shows length-dependent diffusion of AF488-labeled mucin mimetics **GP** in RBC membranes. Lines represent the average signal from n = 6 cells, c_**GP**_ = 7.5 uM), *p-values were determined by one-way ANOVA with multiple comparisons (* < 0.05, ** < 0.01*).

RBCs provide a useful model for examining the effects of glycocalyx properties on lectin and IAV interactions with sialoglycan receptors. RBCs present endogenous Sia modifications in both α2,3 and α2,6 linkages and are commonly used to assess IAV binding activity via cell hemagglutination. Even though this cell type is not the target of IAV infection in mammals, RBCs also offer a useful platform to study mucin-related phenomena by providing an intermediary between simple laboratory membrane models based on supported lipid bilayers and lipid vesicles containing chemically defined glycoconjugates and the dynamic and heterogeneous environment of the epithelial cell glycocalyx. The RBC present compositionally complex but compact native glycocalyx (~ 5-10 nm),^25^ which can be augmented with glycopolymers to introduce extended mucin-like features. At the same time, RBCs lack endocytic activity, which allows for installing mucin-mimetic brushes at high-density and compositional stability.

To introduce mucin-like features into the RBC glycocalyx, the cells were treated with the lipid-terminated glycopolymers **GP** of all three lengths at increasing concentrations (c_**GP**_ = 0.05 – 50.00 μM) for 1 hour. Membrane incorporation of the fluorescently labeled polymers was measured by flow cytometry (**Fig 2C**). The efficiency of membrane insertion was inversely proportional to polymer length, resulting in increasingly extended but less dense glycocalyx structures when the cells were treated at equal polymer concentrations. Polymers lacking the seco-cholestenone hydrophobic anchor did not insert into the cell membranes (**Fig S12**).

Examination of the cells with light microscopy revealed pronounced rounding and loss of membrane features with increasing incorporation of the medium and long polymers (**Fig 2D**, shown for **GP-L**). Flow cytometry analysis of RBCs after treatment with mucin mimetics of all three lengths (c_**GP**_ = 50 μM) showed negative correlation between **GP** length and side scatter (SSC) with little change in forward scatter (FSC) intensity (**Fig S13**). This supports the visual observation of a polymer size-dependent decrease in cell complexity without significant alterations of the overall cell size (**Fig 2D**). The changes in cell morphology likely arise from entropic pressures exerted on the cell membrane upon a mushroom-to-brush conformational transition of the tethered mucin-mimetics in response to glycocalyx crowding, as proposed recently by Paszek and co-workers in their analysis of the role of mucins in cell-shape regulation.^26^ The transition in cell shape was accompanied by changes in sedimentation properties of the remodeled RBCs. We identified polymer concentration thresholds of ~ 7.5 μM and 2.0 μM for **GP-M** and **GP-L**, respectively, above which cell sedimentation becomes less efficient (**Fig 2E**). No such effect was detected for the shortest polymers **GP-S** over the tested range of concentrations. RBC aggregation and sedimentation are processes facilitated by the cells’ discoid shape and surface charge, which are both features affected by the introduction of the extended mucin-like glycocalyx.^27^

The morphological changes in the cells induced by the mucin mimetics indicate an increase in glycocalyx crowding, which should likewise influence the lateral membrane diffusion of the glycopolymers. Taking advantage of the fluorophore labels present in the polymers, we performed fluorescence recovery after photobleaching (FRAP) analysis on cells treated with **GPs** of all three lengths at the threshold concentration, c_**GP**_ = 7.5 μM, at which the medium-size mucin mimetic **GP-M** induced alterations in cell shape and sedimentation properties (**Fig 2F**). This concentration is above the transition concentration for the long polymer **GP-L** and close to the surface saturation level for the short mimetic, **GP-S**, which did not alter sedimentation. The FRAP analysis revealed that all three glycopolymers showed lateral diffusion within the endogenous glycocalyx, with fluorescence recovery rates decreasing from 0.91 ± 0.24 s^−1^ for **GP-S**, to 0.36 ± 0.08 s^−1^ for **GP-M**, to 0.18 ± 0.04 s^−1^ for the longest polymer, **GP-L**. Considering that the cell surface grafting efficiency for **GP-S** is significantly higher than for **GP-M** and **GP-L** (**Fig S11**), the mobility of the mucin mimetics in the glycocalyx is, thus, influenced more strongly by their size rather than their density in the membrane.

### Mucin mimetics slow the rate of lectin association while enhancing binding complex stability

We examined the effects of the mucin-mimetic glycocalyx size and density on the binding of *Sambucus nigra* agglutinin (SNA) to Sias on RBCs. Cells treated with **GPs** of all three lengths near the threshold concentration for inducing cell morphology change (c_**GP**_ = 7.5 μM) were incubated with a fluorescently labeled (AF647) SNA lectin at sub-agglutination concentration (*c_**SNA**_* = 0.2 μg/mL). The lectin binding was assessed via flow cytometry at regular time points until signal saturation (~ 13 min) and the resulting data were fitted and benchmarked to untreated control cells to determine the relative rates (*r_rel_*) of binding for each condition (**Fig 3a**). The medium and long glycopolymers **GP-M** and **GP-L** reduced the rates of SNA binding to a similar extent (*r_rel_* = 0.84 +/− 0.05 and 0.88 +/− 0.05 s^−1^, respectively), while the shortest mucin mimetic, **GP-S**, had no effect on the extent or rate of lectin association (*r_rel_* = 1.02 +/− 0.05). These observations are consistent with glycopolymer shielding of endogenous glycan receptors^18^ according to their size and is expected based on the dimensions of the RBC glycocalyx (~ 10 nm) and the lengths of the different mucin mimetics (~ 3, 13 and 40 nm, **Fig 2a**).

**Figure 3.**
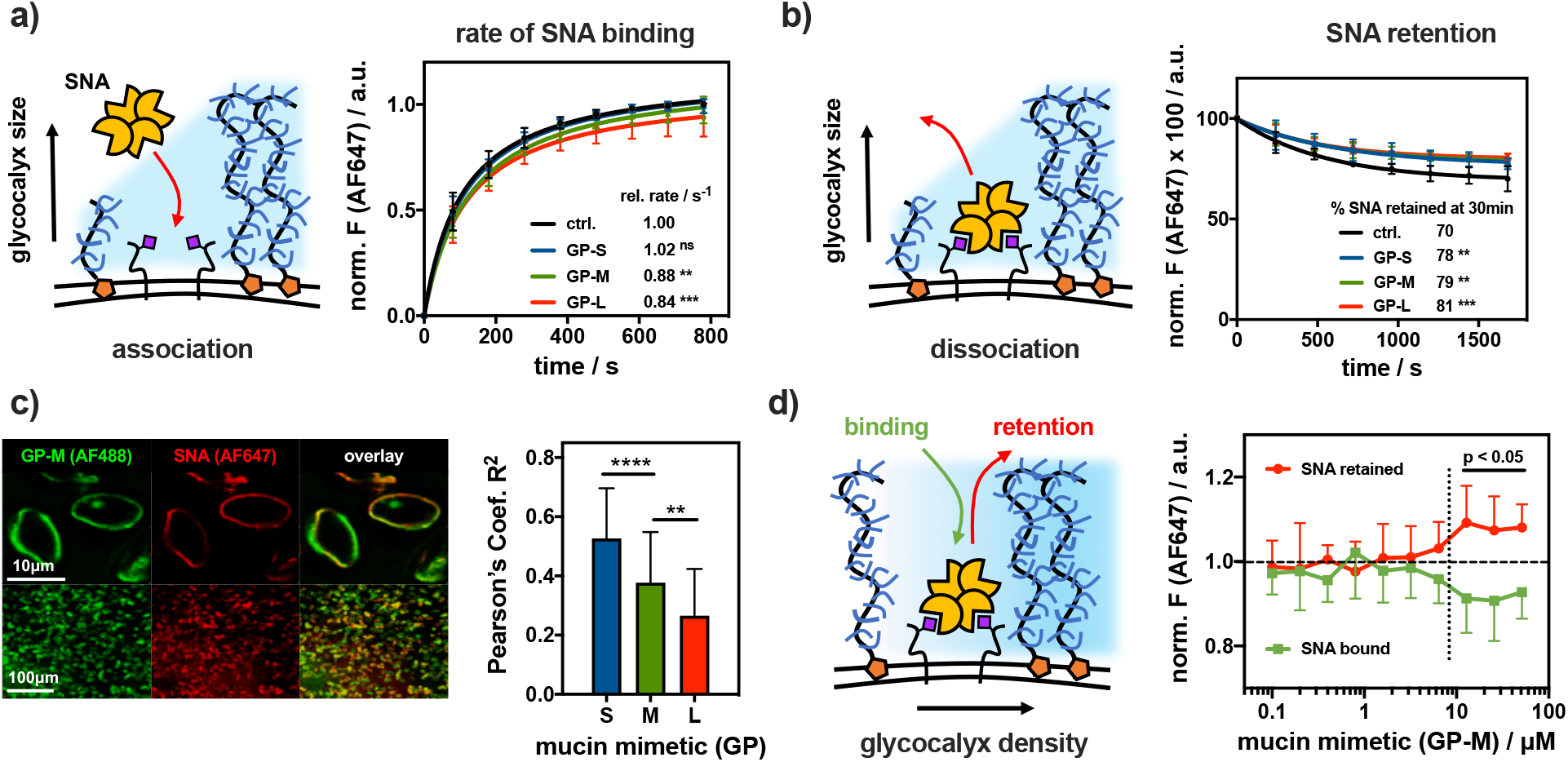
Spectator glycocalyx size and density regulate SNA interactions with cell surface receptors. **a)** Extended glycocalyx reduces the rate of SNA binding. Red blood cells (RBCs) were remodeled with short (**S**), medium (**M**) and long (**L**) mucin mimetics **GP** (c_**GP**_ = 7.5 μM) and the binding for AF647-SNA (c_**SNA**_ = 0.2 μg/mL) was measured by flow cytometry and normalized to untreated cells. (n=6 experimental replicates). **b)** Extended glycocalyx reduces SNA dissociation from RBCs. The remodeled cells equilibrated AF647-SNA were resuspended in pure buffer and lectin retention was measured by flow cytometry. (n=6 experimental replicates). **c)** Extended mucin mimetic glycocalyx drives segregation of SNA-receptor adhesion complexes. Florescence micrographs show representative confocal images of RBCs remodeled with mucin mimetic **GP-M** (green, c_**GP**_ = 7.5 μM) and stained with AF647-SNA (red, c_**SNA**_ = 0.2 μg/mL). Bar graph represents Pearson’s correlation coefficient (R^2^) analysis of images for SNA binding to RBCs remodeled with polymers of all three lengths (c_**GP**_ = 7.5 μM). (n > 35 individual cell images per polymer condition). **d)** Glycocalyx crowding limits SNA association with cell surface receptors while stabilizing the resulting adhesion complex. Association (green) and retention (red) of AF647-SNA at the surface of RBCs remodeled with **GP-M** at increasing concentration were measured by flow cytometry and normalized to untreated cells. (n=12 experimental replicates). *Values represent averages and standard deviations, p-values were determined by ANOVA (a-c) or student test (d), *<0.05, **<0.01, ***<0.001, ****<0.0001.*

To assess the effects of the mucin mimetics on lectin retention at the cell surface, the remodeled RBC were allowed to reach saturation binding with SNA (15 min), washed, and allowed to re-equilibrate in fresh PBS buffer (30 min). Flow cytometry was used to monitor the retention of the fluorescent SNA by RBCs over the course of the experiment and the percentage of SNA remaining at equilibrium was calculated (**Fig 3b**). Polymers of all three lengths enhanced SNA retention by ~ 8-11% over untreated control cells. In contrast to lectin association, the impact of polymer length on SNA retention was much less pronounced, with the shortest polymer **GP-S** also exerting significant effect. This points to a change in the avidity of the SNA-receptor binding complex driven by polymer density rather than size. Fluorescence confocal microscopy showed increased exclusion of the mucin mimetics from SNA (**Fig 3c,** c_**GP**_ = 7.5 μM, c_**SNA**_ = 0.2 μg/mL), suggesting extensive lectin clustering in the presence of the polymers. Pearson’s correlation analysis (R^2^ values) showed a decrease in glycopolymer colocalization with SNA according to size (**GP-S**, R^2^ = 0.53 ± 0.17; **GP-M**, R^2^ = 0.38 ± 0.17; **GP-L**, R^2^ = 0.27 ± 0.16).

Our experiments indicate that increasing mucin mimetic size inhibits lectin association, their membrane crowding promotes the formation of more stable adhesion complexes, presumably through lectin clustering. In the mucin glycocalyx, which can be described as a surface-tethered polymer brush,^26^ the extension of the glycoproteins away from the membrane can be driven by crowding. Similar behavior was also recently confirmed for synthetic glycopolymers in supported lipid bilayers.^18^ To evaluate the effect of mucin-mimetic density in the RBC glycocalyx on SNA interactions, we determined equilibrium binding and retention of SNA at increasing membrane densities for the medium-sized polymer **GP-M** (c_**GP-M**_ = 0.1 – 50.0 μM). We observed an emergence of SNA shielding at polymer concentration of 6.4 μM, which also coincided with the transition of the RBCs toward a more rounded phenotype with altered sedimentation properties (**Fig 2E**). Likewise, SNA dissociation became significantly inhibited at this threshold concentration (**Fig 3b**). Together, these observations support a density-dependent transition of the glycopolymers from coiled to extended conformations in the glycocalyx brush, as predicted by the current biophysical model for the behavior of membrane-associated mucins.^26^ The shielding ability of the mucin-mimetics was further enhanced at higher concentrations of SNA (**Fig S16**), indicating that the lectin also contributes to glycocalyx crowding once bound to its target receptors.

### Mucin mimetics drive lectin clustering by increasing glycocalyx crowding

The low levels of colocalization between the mucin mimetics and SNA in the RBC glycocalyx suggest increased clustering of the oligomeric lectin, which may stabilize the binding complex through crosslinking of neighboring sialylated glycoconjugates in the plasma membrane. To confirm enhanced lectin clustering by the glycopolymers, we measured Förster resonance energy transfer (FRET) between SNA probes labeled separately with AF594 (donor) or AF647 (acceptor) fluorophores by fluorescence lifetime imaging microscopy (FLIM). A shortening of the donor excited state lifetime (τ) indicates increased FRET efficiency, and thus reduced distance, between the donor and acceptor fluorophores.

First, RBCs were treated with the SNA labeled with AF594 only for 15 min and imaged by FLIM to establish the fluorescence lifetime of the donor (τ = 3.15 ± 0.02 ns) (**Fig 4a,** c_**SNA**_ = 0.2 μg/mL). Incubation of the cells with equimolar amounts of the donor AF594-SNA and acceptor AF647-SNA probes reduced the donor excited state lifetime (τ = 2.53 ± 0.09 ns), setting the baseline for SNA proximity in the RBC glycocalyx in the absence of the mucin mimetics. Repeating this experiment in the presence of the long glycopolymer **GP-L** (c_**GP**_ = 7.5 μM) further reduced the donor excited state lifetime (τ = 1.80 ± 0.07 ns), revealing closer spacing of the bound lectins. Control experiments with cells treated with the *seco*-cholestenone anchor only (c_**chol**_ = 7.5 μM) did not reduce lectin spacing (τ = 2.65 ± 0.21 ns) compared to RBC controls without mucin mimetic treatment, confirming that the glycopolymer domains were responsible for driving lectin clustering.

**Figure 4.**
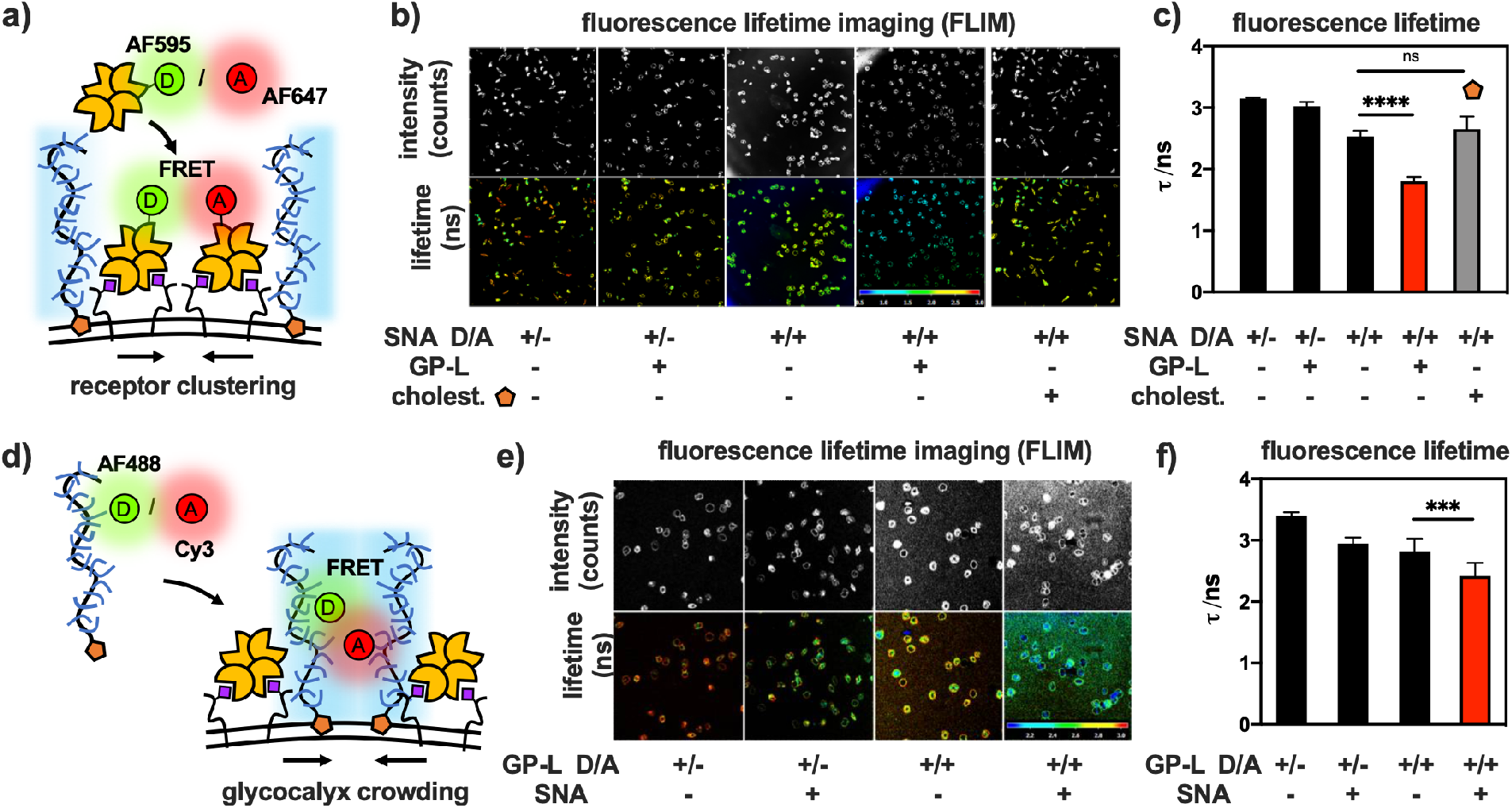
Glycocalyx crowding drives clustering of lectin-receptor adhesion complexes. **a**) Changes in SNA clustering in response to glycocalyx crowding were assessed based on Förster resonance energy transfer (FRET) between lectins labeled with donor (D = AF595) and acceptor (A = AF647) probes. **b**) Representative fluorescence lifetime imaging microscopy (FLIM) images and **c**) bar graph representations for the binding of SNA probes to RBCs before and after remodeling with **GP-L** (c_**GP**_ = 7.5 μM) or the hydrophobic anchor **S.5** alone (c_**S.5**_ = 7.5 μM). Decrease in fluorescence lifetime (t) indicates closer SNA proximity. **d)** Mucin mimetics labeled with donor (D = AF488) an acceptor (A = Cy3) probes were employed to measure changes in glycocalyx crowding after SNA binding via FRET. **e**) Representative FLIM images and **f**) bar graph representations of RBCs remodeled with mucin-mimetic probes (c_**GP-D/A**_ = 15 μM) before and after incubation with SNA. Decrease in fluorescence lifetime (t) indicates enhanced glycocalyx crowding. Color scales represent fluorescence lifetime (t/ns). *Values represent averages and standard deviations for representative image frames containing >10 cells, p-values were determined by student test, ***<0.001, ****<0.0001.*

In the absence of ligand-dependent molecular interactions between the glycopolymers and SNA, the observed clustering of the lectin is likely induced by the biophysical properties and membrane diffusion of the mucin mimetics. Enhanced expression of mucins has been shown to increase glycocalyx crowding, which can manifest through changes in membrane shape and the clustering of cell adhesion complexes.^26,16^ We used FRET/FLIM to analyze the crowding of mucin mimetics in the RBC glycocalyx in response to SNA binding by measuring changes in their proximity (**Fig 4b**). Long glycopolymers **GP-L** with identical size and glycan composition were synthesized to display donor (AF488) or acceptor (Cy3) fluorophores with overlapping emission and absorption profiles to enable FRET. RBCs were treated either with polymers containing the donor fluorophore only or with an equimolar mixture of polymers presenting both the donor and acceptor dyes (total c_**GP**_ = 15μM). FLIM imaging of RBCs remodeled with the glycopolymer FRET pair showed significantly reduced donor fluorophore lifetime from τ = 2.95 ± 0.10 ns to τ = 2.42 ± 0.21 ns after incubation with SNA (c_**SNA**_ = 0.2 μg/mL) for 15 min. The higher FRET efficiency due to shorter distances between the mucin mimetic probes provides experimental support for increased crowding of the glycocalyx environment in response to SNA binding.

Collectively, the FRET/FLIM experiments showed that the lectin clustering and mucin mimetic crowding phenomena are interconnected and occur simultaneously. If the mucin mimetics forced changes in receptor organization prior to lectin binding, we would not except to observe significant changes in FRET efficiency from the polymers after introduction of SNA. FRAP analysis shows that, once binding equilibrium is reached, the lectins become immobile in the RBC glycocalyx (**Fig S17**), providing further support for the stabilization of the SNA binding complexes through clustering and receptor crosslinking.

### Glycocalyx crowding with mucin mimetics enhances H1N1 virus retention on RBCs

We sought to evaluate whether the paradox observed for the effect of mucin mimetics on SNA binding, whereby the extended glycopolymers slowed the kinetics of lectin association while stabilizing the resulting binding complex, would also apply to viruses that utilize multivalent receptor interactions for adhesion to host cells.

First, we measured the effects of mucin mimetic length and membrane density on the adhesion of the H1N1 virus to RBCs. Cells remodeled with mucin mimetic glycopolymers **GPs** of all three lengths (c_**GP**_ = 7.5 μM) alongside untreated controls were incubated with H1N1 (33 HAU) for 15 min. The binding experiments were performed at room temperature to limit neuraminidase (NA) activity and viral release from the cell surface. After incubation, H1N1 binding was detected using a NA enzymatic assay with the fluorogenic reporter substrate, 4-methylumbelliferyl *N*-acetyl-α-neuraminic acid (4-MU-NANA) at 37°C.^28^ The rate of fluorescence signal generation by neuraminidase activity was used to determine the concentration of RBC-bound viruses in the sample (**Fig S18**). All three mucin mimetics reduced the overall amount of virus bound to the RBCs compared to non-treated cells (**Fig 5a**). The efficiency of viral binding inhibition correlated with polymer length, increasing from ~ 20% for the short mimetic **GP-S** to ~ 50% for the long polymer **GP-L**, consistent with enhanced shielding of cell-surface receptors by the progressively extended glycocalyx. The shielding capacity of the mucin mimetic glycocalyx against viral adhesion increased with membrane density of the polymers and became most effective at the threshold polymer concentration associated with changes in RBC shape and sedimentation properties (**Fig 5b**, c_**GP-M**_ = 6.3 μM). As observed for SNA, increasing viral titers reinforced the protective properties of the mucin mimetics, presumably, by further increasing local crowding in the glycocalyx (**Fig S19**).

**Figure 5.**
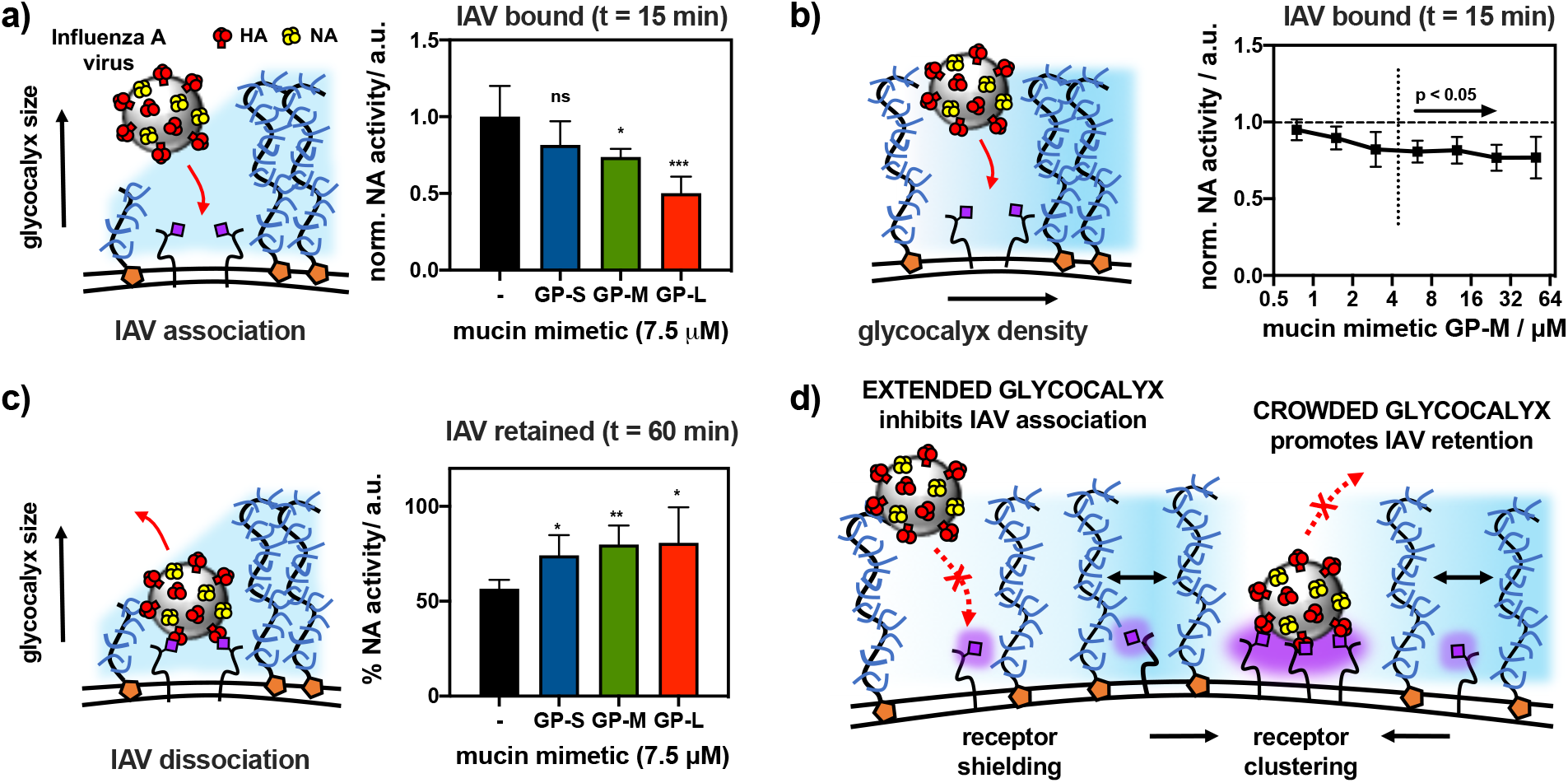
Influence of glycocalyx size and density on the binding of H1N1 viruses to sialic acid receptors on RBCs. **a)** Extended glycocalyx shields endogenous sialic acid receptors from H1N1 binding. Saturation binding of H1N1 (t = 15 min) to RBCs remodeled with short (**S**), medium (**M**) and long (**L**) mucin mimetics **GP** (c_**GP**_ = 7.5 μM) was assessed via their neuraminidase (NA) activity toward a fluorogenic substrate 4MU-NANA. **b)** Increased glycocalyx crowding limits viral adhesion. Saturation binding of H1N1 (t = 15 min) to RBCs remodeled with increasing concentrations of **GP-M** (c_**GP**_ = 0.8 - 50.0 μM) was evaluated via NA activity assay with 4MU-NANA. **c)** Extended glycocalyx enhances retention of viruses bound to RBC receptors. Retention of H1N1 viruses by RBCs remodeled with mucin mimetics **GP** of increasing length (c_**GP**_ = 7.5 μM) was measured as a fraction of NA activity before and after equilibration in fresh buffer (t = 60 min). **d)** Proposed model for the influence of a spectator mucin-type glycocalyx on cell-pathogen interactions. Extended and dense glycocalyx shields cell surface receptors from viral adhesion. Glycocalyx crowding drives receptor clustering and enhances retention of bound viruses. *Values represent averages and standard deviations for n = 6 experimental replicates; p-values were determined by t-test (a and c) and ANOVA (b): *<0.05, **<0.01, ***<0.001, ****<0.0001.*

Analysis of H1N1 retention on RBCs confirmed the ability of the mucin mimetic glycocalyx to stabilize the adhesion complex between the virus and its sialoglycan receptors at the cell surface (**Fig 5c**). RBCs remodeled with glycopolymers of all three lengths (c_**GP**_ = 7.5 μM) as well as untreated control cells were first allowed to equilibrate with H1N1 (33 HAU) for 30 minutes, then they were washed to remove any free virus and resuspended in free buffer for 60 min to allow for dissociation. The experiments were carried out at room temperature to minimize NA activity. The levels of virus bound to the RBC surface before and after dissociation were measured via the 4-MU-NANA fluorescence NA activity assay as described above. Mucin mimetic polymers of all three lengths increased the amount of H1N1 that was retained at the cell surface (75-80%) compared to untreated RBC controls (57 %). As observed for SNA, the effect of polymer size on enhancing H1N1 retention was less prominent compared to the inhibition of viral binding (**Fig 3 and Fig 5a**), pointing to a polymer density-driven stabilization of the adhesion complex.

## Discussion

Membrane-associated mucins contribute to the protection of epithelial surfaces against pathogen invasion in two distinct ways. Mucins present decoy receptors for capturing pathogens and shedding them from the cell surface. Their organization into a dense extended brush also presents a physical barrier blocking the pathogen from accessing functional receptors at the cell surface. Examining how each mechanism contributes to the protection of host cells against pathogen invasion may guide the development of new therapeutics targeting early stages of infection.

While a range of techniques exist to profile the glycan-binding specificity of pathogen-associated adhesion proteins, the repertoire of tools to study the biophysical properties of the mucin glycocalyx remains limited. Recently, genetic tools have begun to emerge that enable expression of recombinant mucins with precisely defined lengths and membrane densities in cells. This has provided new insights into how the composition and physical properties of the mucin glycocalyx contribute to membrane shape generation^26^ and modulation of cell adhesion.^29^ Although the structure of mucins can now be tailored, tuning their glycosylation state (e.g., the removal or addition of sialic acids) without affecting other glycoproteins and glycolipids at the cell surface remains a challenge. This makes decoupling the decoy receptor and physical barrier functions of membrane-associated mucins in protecting cells against pathogens, particularly in instances, when both protective mechanisms are deployed simultaneously.

One example where membrane mucins serve both purposes is in preventing IAV adhesion to host cells. The mucins are heavily sialylated and capture IAVs by binding to the viral HA proteins. And while the Sia decoys are rapidly removed by viral NAs, the mucins remain on the cell surface and continue to pose a physical barrier against the virus. Genetic studies identified mucin MUC1 as an important factor in limiting H1N1 infection by shedding the bound virus from the cell surface after the mucin is proteolytically cleaved.^11^ Although the inhibition of IAV binding by physically shielding non-mucin Sia receptors on the cell surface is yet to be demonstrated, studies with *H. pylori* lacking the mucin adhesins, BabA and SabA, showed inhibition of bacterial infection with increased MUC1 expression.^10^

To examine the biophysical principles through which membrane-associated mucins limit pathogen association with host cells, we have generated artificial glycocalyx models of the mucin like brush on the surfaces of RBCs, which present a biologically relevant environment with respect to receptor complexity and heterogeneity. Using synthetic glycopolymers with tunable size displaying non-participating model glycans and endowed with hydrophobic plasma membrane anchors and fluorescent optical probes, we were able to tailor the features of the mucin-mimetic brush with respect to size and density and characterize its dynamics. Using our RBC-based mucin glycocalyx model, we systematically evaluated how the length and membrane density of the mucin mimetics influences both binding and dissociation of SNA and H1N1 at the cell surface. By selecting glucose as the side chain glycan residue for our mimetics, we ensured that the probes would not interact with SNA, H1N1, or any other endogenous glycan-binding protein on the RBC surface. Therefore, any observed changes in lectin or viral binding could be ascribed solely to the biophysical properties of the glycocalyx component.

Installation of the mucin mimetics on the RBC surface introduced significant glycocalyx crowding, which manifested by changes in cell shape toward more rounded morphology, consistent with recent studies showing that increased expression of mucins by cells enforced membrane curvature.^26^ The change in morphology was accompanied by altered sedimentation properties of the cells, indicating changes in the physical properties of the glycocalyx. FRAP experiments confirmed that the polymers retained their mobility in the glycocalyx through lateral diffusion in the membrane and the rate of diffusion was inversely correlated to polymer length, indicating physical interactions of the extra-membrane glycopolymer domain with endogenous glycocalyx components. Analysis of SNA and H1N1 binding near the threshold polymer density associated with morphological and sedimentation changes in the RBCs slowed the association rate of the lectin by ~ 12-16 % and reduced the overall amount of virus captured at the cell surface by 20-50%. The protective effects of the mucin mimetics were polymer length- and membrane density-dependent, in agreement with the proposed mucin brush shielding model and the recent study by Bertozzi and coworkers reporting inhibition of H1N1 binding to gangliosides by mucin-mimetic brushes based on lactosylated poly-L-serine polymers anchored in supported lipid bilayers (SLBs).^18^

Unexpectedly, the mucin mimetics enhanced the retention of both SNA (8-11%) and H1N1 (18-23%) at the RBC surface compared to untreated cell controls. This effect was independent of glycopolymer size and appeared to be driven primarily by glycocalyx crowding. This suggests that glycan-binding proteins and pathogens may exploit the crowded glycocalyx interface of the epithelial mucosa to strengthen their association with host cells. Confocal fluorescence microscopy of RBC stained with SNA revealed increased exclusion of the mucin mimetics from the lectin adhesion sites based on their length. FLIM/FRET imaging further showed more extensive clustering of SNA in the presence of the mucin mimetics as well as increased crowding in the excluded mucin brush upon lectin binding.

Based on our observations, we propose a biophysical mechanism for strengthening of interactions between soluble and virus-associated oligomeric lectins and cell surface glycan receptors (**Fig 5d**). Therein, the binding of the protein to its receptors induces exclusion of the mucin-mimetics from the point of contact and induces local crowding in the glycocalyx. To alleviate the crowding developing in the mucin mimetic brush,^26^ smaller glycoconjugates may diffuse back toward the adhesion complex. If these glycoproteins and glycolipids contain Sia modifications, this would generate a higher-valency adhesion patch increasing lectin clustering and overall avidity of the binding event. The limited effect of polymer size on SNA and H1N1 retention supports the proposed mechanism, as the density of mucins at the cell surface, not their length, has been previously identified as the primary driver of entropic pressure on the cell membrane influencing its shape and stability.^26^ The rounding of RBCs with increasing mucin mimetic density at their membranes provide further evidence for glycocalyx crowding. The proposed mechanism does not require direct binding of the pathogenic adhesins to mucins but assumes multivalency of their interactions with glycan receptors in the glycocalyx. Since most glycan binding proteins utilize multivalency to achieve avidity, we anticipate that this mechanism will be general and exploited by other pathogens that target glycan receptors in mucosal tissues.

## Conclusion

In this study, we have applied cell surface engineering with synthetic mimetics of mucin glycoproteins to model the mucosal glycocalyx. By tuning the nanoscale dimension and membrane density of the materials, we assessed the effectiveness of the physical barrier to restrict lectin and virus adhesion to target receptors. The extended and dense cell surface displays of mucin mimetics inhibited the rate of SNA and H1N1 binding, consistent with the current view of mucins acting as a physical shield protecting the cell surface. We discovered that the both the soluble and virus-associated oligomeric lectins exploited the crowded dynamic glycocalyx interface to strengthen their interactions with cell-surface receptors. The mucin mimetics were designed to be refractory to the tested glycan-targeting proteins, as such the observed behavior was driven by the biophysical properties of the glycocalyx and may be a general feature of the mucosal barrier.

## Acknowledgements

We thank the UCSD Microscopy Core facility (via p30 grant NS047101 from NINDS) for assistance with fluorescence microscopy imaging, and the Glycobiology Research and Training Center for access to tissue culture facilities and analytical instrumentation. The authors acknowledge the use of facilities and instrumentation supported by the NSF through the UC San Diego Materials Research Science and Engineering Center (UCSD MRSEC), DMR-2011924. We also wish to thank Dr. Christopher Fisher for help with H1N1 production and Taryn Lucas for her assistance with neuraminidase assays. This work was supported by the NIH Director’s New Innovator Award (NICHD: 1DP2HD087954-01). K. G. is supported by the Alfred P. Sloan Foundation (FG-2017-9094) and the Research Corporation for Science Advancement via the Cottrell Scholar Award (grant # 24119).

