## Supporting information for "Glycocalyx crowding with synthetic mucin mimetics strengthens interactions between soluble and virus-associated lectins and cell surface glycan receptors"

### Instrumentation and reagents.

**Instrumentation.** Column chromatography was performed on a Biotage Isolera One automated flash chromatography system. Nuclear magnetic resonance (NMR) spectra were collected on a Bruker 300 MHz and a Jeol 500 MHz NMR spectrometers. Spectra were recorded in $CDCl_3$ or $D_2O$ solutions at 293K and are reported in parts per million (ppm) on the δ scale relative to the residual solvent as an internal standard (for $^1H$ NMR: $CDCl_3$ = 7.26 ppm, $D_2O$ = 4.79 ppm, for $^{13}C$ NMR: $CDCl_3$ = 77.0 ppm). HRMS (high-resolution mass spectrometry) analysis was performed on an Agilent 6230 ESI-TOFMS in positive ion mode. UV-Vis spectra for polymer fluorophore content quantification were collected using a quartz cuvette using a Thermo Scientific Nanodrop2000c spectrophotometer. UV-Vis spectra for kinetic measurement of 4MU-NANA fluorescence turn on was collected in 96 well plate format using a SpectraMax i3x (Molecular Devices). IR spectroscopy was performed on a Nicolet 6700 FT-IR spectrophotometer (Thermo Scientific). Size exclusion chromatography (SEC) was performed on a Hitachi Chromaster system equipped with an RI detector and two 5 µm, mixed bed, 7.8 mm I.D. x 30 cm TSKgel columns in series (Tosoh Bioscience). Organic soluble polymers were analyzed using an isocratic method with a flow rate of 0.7 mL/min in DMF (0.2% LiBr, 70 °C). For aqueous SEC, two 8 µm, mixed-M bed, 7.5 mm I.D. x 30 cm PL aquagel-OH columns in series (Agilent Technologies) were run in sequence using an isocratic method with a flow rate of 1.0mL/min in water (0.2M $NaNO_3$ in 0.01M $Na_2HPO_4$, pH = 7.0). Flow cytometry analysis was performed on live RBCs using a FACS Canto II cytometer (BD Biosciences). Microscopy techniques were performed on either a Keyence Fluorescent microscope (brightfield) or Leica SP5 (all fluorescence techniques).

**Materials.** All chemicals, unless stated otherwise, were purchased from Sigma Aldrich and used as received. Reaction progress was checked by analytical thin-layer chromatography (TLC, Merck silica gel 60 F-254 plates) monitored either with UV illumination, or by staining with iodine, ninhydrin, or CAM stain. Solvent compositions are reported on a volume/volume (v/v) basis unless otherwise noted. 4,5-seco-cholesten-5-one[1] and Glc[2] propargyl glycosides were prepared according to published procedures. Turkey Red Blood Cells as a 10% solution were obtained from Lampire Biological Laboratories (cat # 724908). *Sambucus nigra* agglutinin (SNA) lectin were purchased from Vector Labs. NHS functionalized AlexaFluor 594 (AF 594) and AlexaFluor 647 (AF 647) for lectin labeling were purchased from Sigma Aldrich, and AlexaFluor 488 (AF 488) alkyne Cyanine 3 (Cy3)-alkyne for labeling of polymers was obtained from Sigma Aldrich. The sialidase reporter molecule 2'-(4-methylumbelliferyl)-α-D-N-acetylneuraminic acid (4MU-NANA) was purchased from Carbosynth. β-propargyl glucoside (Glc) was prepared according to a previously published procedure.[3]

### Synthesis of glycopolymers GP-S/M/L.

**Scheme S1.** Synthesis of Glycopolymers through iterative CuAAc click strategy.

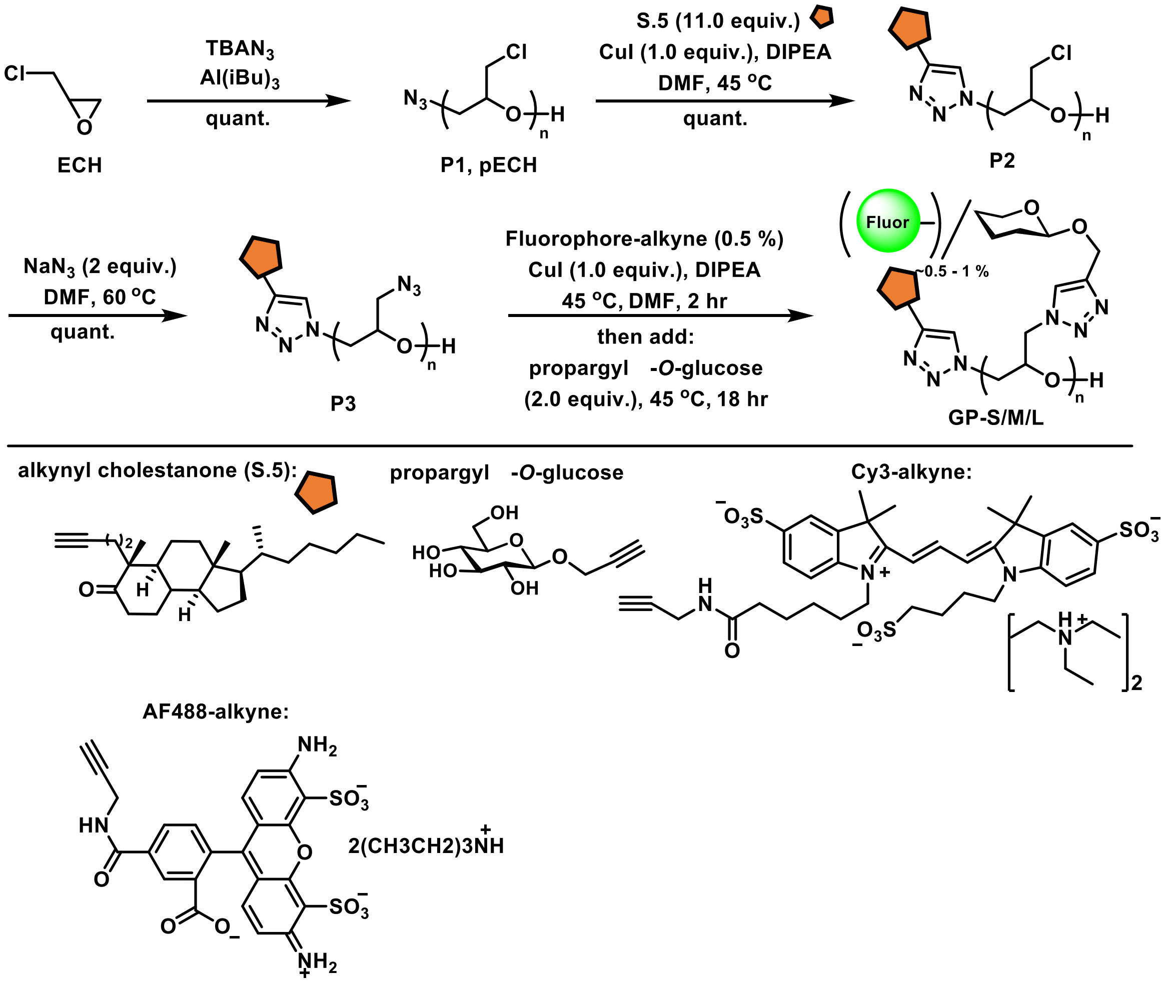

#### General Procedure for the preparation of poly(epichlorohydrin) (P1).

Epichlorohydrin was polymerized according to the procedure developed by Carlotti.[4] Briefly, for the longest glycopolymers **GP-L** as a representative example, a flame-dried Schlenk flask (10 mL) equipped with a magnetic stirrer and fitted with a PTFE stopcock was charged with tetrabutylammonium azide (TBAN3, 20 mg, 0.037 mmol, 0.002 equiv.) under argon. A solution of freshly distilled epichlorohydrin (1.29 mL, 16.5 mmol) was prepared in anhydrous toluene (4 mL). A solution of triisobutylaluminum in toluene (1.07 M, 104 μL, 0.111 mmol, 0.007 equiv.) was added via a syringe under argon at −30 °C. The reaction was stirred for 4 hours and then stopped by the addition of ethanol. The resulting pECH polymer 1 was precipitated into hexanes and dried under vacuum to yield a clear viscous oil (1500 mg, 99% yield). The polymer was analyzed by SEC (0.2% LiBr in DMF), IR, and NMR.

**General Procedure for the end-functionalization of poly(epichlorohydrin) (P2).**

For the longest glycopolymers **GP-L** as a representative example, in a flame-dried Schlenk flask (10 mL), pECH polymer **P1** (15 mg, 0.3 μmol) was dissolved in degassed anhydrous DMSO (200 μL). 4,5-seco-cholesten-5-one **S.5** (1.7 mg, 3.8 μmol, 11.0 equiv.) was added, followed by CuI (~0.05 mg, 0.3 μmol, 1.0 equiv.) and one drop diisopropylethyl amine (DIPEA, ~5 μL). The reaction was stirred at 40 °C for 12 h, at which time it was quenched by the addition of water to precipitate the polymer. The resultant polymer was triturated with hexanes to remove unreacted **S.5** and dried on vacuum to yield a clear viscous oil in (16 mg, 100% yield). The polymer was analyzed by SEC (0.2% LiBr in DMF).

**General Procedure for the preparation of poly(Glycidyl Azide) (P3).**

The chloride to azide exchange in pECH polymer **P3** was accomplished according to a previously published procedure.[5] Briefly, in a flame-dried Schlenk flask (10 mL), polymer **P2** (15 mg, 0.16 mmol) was dissolved in dry DMF (300 μL). To the solution was added $NaN_3$ (21 mg, 0.32 mmol, 2.0 equiv.), and the reaction was stirred at 60 °C for 3 days under argon to allow complete conversion. The polymer solution was filtered and precipitated in ethanol to yield a clear viscous oil (16 mg, 100% yield). The polymer was analyzed by SEC (0.2% LiBr in DMF.

**General procedure for the preparation of Glycopolymers (GP-S/M/L).**

In a flame-dried Schlenk flask (10 mL), polymer **P3** (9.00 mg, 0.09 mmol) was dissolved in degassed dry DMSO (250 μL). To the solution was added AF488-alkyne or Cy3-alkyne (0.50 mg, 0.50 μmol) in DMSO (50 μL), followed by CuI (2.00 mg, 9.00 μmol) and DIPEA (16 μL, 0.09

mmol). The reaction was stirred in the dark under Ar at 40 °C for 2 h. After this time, β-propargyl glucoside (0.02 mmol, 1.50 equiv. per azide side-chain) in degassed anhydrous DMSO (50 μL) was added to the reaction mixture. The reactions were stirred in the dark at 40 °C overnight. After this time, the reactions were diluted with DI water and treated with Cuprisorb beads (SeaChem labs) for 18 h to sequester copper. The resulting copper-free solutions were filtered through Celite to remove the resin and lyophilized. The dry residues were triturated 3× with methanol with monitoring by TLC to remove excess unreacted glycosides. The resulting AF488 or Cy3-labeled glycopolymers **GP-S/M/L** were dissolved in D2O and lyophilized to give a blue solid in a quantitative yield for each polymer. The polymers were characterized using 1H NMR (D2O, 300 MHz), IR, and UV-Vis (λmax = 488 or 554 nm) spectroscopy. Absorbance readings at known concentrations of glycopolymers **GP-S/M/L** indicated the presence of 0.3 – 2.7 fluorophores per polymer chain depending upon polymer length (~0.5 - 1% sidechain occupancy). The theoretical Mn of the final glycopolymers **GP** were calculated assuming 100% sidechain substitution with glucose.

### Methods for biological assays.

#### Remodeling of RBC glycocalyx with glycopolymers GP.

RBCs (4% w/v in PBS) were incubated with AF488-labeled glycopolymers **GP-S/M/L** at increasing concentrations ($c_{pol}$ = 0.1-30.0 μM) for 1 h at 37 °C. The cells washed 1x with PBS, then were probed for the presence of AF488 fluorescence using flow cytometry. The data was analyzed on Cytobank online software. Cells were gated to exclude debris, and the median fluorescence intensities (MFI) of cells are reported.

#### Sedimentation of glycocalyx-remodeled RBCs.

RBCs (50μL, 1% in PBS) treated with increasing concentrations of glycopolymers **GP-S/M/L** ($c_{pol}$ = 0.1-30 μM) or left untreated were transferred to round-bottom 96 well plates. The cells were allowed to sediment over 45min. After this time, the plates were scanned on an EPSON Perfection V700 Photo scanner (Digital ICE technologies), and the lowest polymer concentrations required to induce RBC agglutination were determined.

**Preparation of RBCs for microscopy.**

RBCs (50uL, 1% in PBS) treated with increasing concentrations of glycopolymers **GP-S/M/L** ($c_{pol}$ = 1-50 μM) or left untreated were diluted 50x and transferred to poly(lysine) slides. The cells were allowed to settle to the slide surface, and excess PBS was removed via pipette prior to application of a cover slip in order to prevent aggregation of cells. Bright field images of RBCs were taken on Keyence fluorescent microscope, and all fluorescence images were taken on a Leica SP5 confocal microscope.

**Determination of membrane fluidity (FRAP).**

FRAP experiments were performed on a Leica SP5 confocal microscope with a 40× water objective. The membranes of RBCs (50uL, 1% in PBS) treated with glycopolymers **GP-S/M/L** ($c_{pol}$ = 7.5 μM) were bleached with a circular spot of diameter ~ 0.5 μm at 488nm wavelength. A single iteration was used for the bleach pulse, and fluorescence recovery was monitored at low laser intensity in 0.11 s intervals for 12 seconds. FRAP was performed on 6 separate cells and then averaged to generate a single FRAP curve. The rate was calculated by fitting the recovery curve to a hyperbola fit in Prism and solving for the rate in the integrated rate equation ($A = A_o$ e^(-kt)).

**Determination of SNA association and dissociation with glycocalyx-remodeled RBCs by flow cytometry.**

In a 96 well round bottom plate, to RBCs (0.33% in PBS) treated with glycopolymers **GP** ($c_{pol}$ =7.5 μM) or 4,5-seco-cholesten-5-one ($c_{chol}$ =7.5 μM), or to untreated cells, were added AF647-labeled SNA lectins at sub-agglutination concentrations ($c_{SNA}$ = 0.2 μg/mL). The cells were vortexed vigorously for ~ 10 s and then analyzed by flow cytometry (Canto II, BD Biosciences) for the presence of AF647 signal at discrete time points until saturation lectin binding was observed. The data were analyzed on Cytobank software. Cells were gated to exclude debris, and median fluorescence intensities (MFI) of cells are reported. Means and standard deviations were calculated from six independent biological experiments, and *p*-values corresponding to each condition vs. untreated RBC control were calculated using ANNOVA tests with PRISM software. The slopes designating the initial rates of lectin association were extracted for each condition and their significance with respect to untreated RBC controls was assessed based on *p*-values calculated using 1-way ANNOVA tests.

**Determination of co-localization between polymer and lectin via fluorescence microscopy.**

RBCs remodeled with glycopolymer ($c_{pol}$=7.5 μM) were added to poly(lysine) slides. Once settled on the surface, AF 647 labeled SNA in PBS ($c_{SNA}$ = 0.2 μg/mL) was incubated on the slide for 15 min. Unreacted lectin was washed away prior to imaging on Leica sp5 confocal microscope. Colocalization was quantified from the average of 12 replicates using Velocity software (Quorum Technologies).

**Determination of polymer and lectin clustering by FLIM.**

RBCs (0.33% in PBS) treated with glycopolymers **GP-L** labeled with AF488, Cy3, or an equimolar mixture of the two ($c_{pol}$ =7.5 μM), or 4,5-seco-cholesten-5-one ($c_{chol}$ =7.5 μM), or untreated cells, were added to poly(lysine) slides. Once settled on the surface, SNA lectins labeled with AF595, AF647, or an equimolar ratio of the two in PBS ($c_{SNA}$ = 0.2 μg/mL) were applied to RBCs on the slide for 15 min. Unreacted lectin was washed away prior to imaging on Leica sp5 confocal microscope. FLIM images were analyzed directly on Leica acquisition software.

**Maintenance of viral culture.**

Influenza virus strain A/PR/8/34 (H1N1, ATCC VR-1469) was purchased from ATCC and propagated in MDCK cells that were transferred to DMEM medium supplemented with 0.2% BSA fraction V, 25mM HEPES buffer, 2 μg/ml TPCK-trypsin, and 1% penicillin/streptomycin ("DMEM-TPCK" media). Viral titers were determined via the hemagglutination test (HAU) using a 1% solution of turkey red blood cells purchased from Lampire.

**Determination of viral binding to remodeled RBCs (4MU-NANA).**

Viral concentration was measured using 4-Methylumbelliferyl N-acetyl-a-D-neuraminic acid (4MU-NANA) fluorescence turn-on reporter for NA activity. Virus (30 HAU) was incubated with 1% RBCs in PBS remodeled with glycopolymers **GP-S/M/L** ($c_{pol}$ = 7.5 μM), or unmodified cells for 15 min. RBCs were centrifuged and unbound virus was removed via pipette. To measure association, cells were then resuspended in 1.2 mM 4MU-NANA and cleavage was monitored by UV/Vis (excitation = 380nm, emission = 450nm) every 60 seconds for one hour at 37 **°C**. To measure retention, cells were instead resuspended in fresh buffer and allowed to equilibrate for 1hr before they were centrifuged, shed virus was removed by pipette, and RBCs were resuspended in 1.2 mM 4MU-NANA and monitored for cleavage as above.

### $^{1}$H NMR spectra of GP-S/M/L and synthetic intermediates.

**Figure S4**. $^{1}$H NMR (300 MHz, $CDCl_3$) of pECH polymers **P1-S/M/L**.

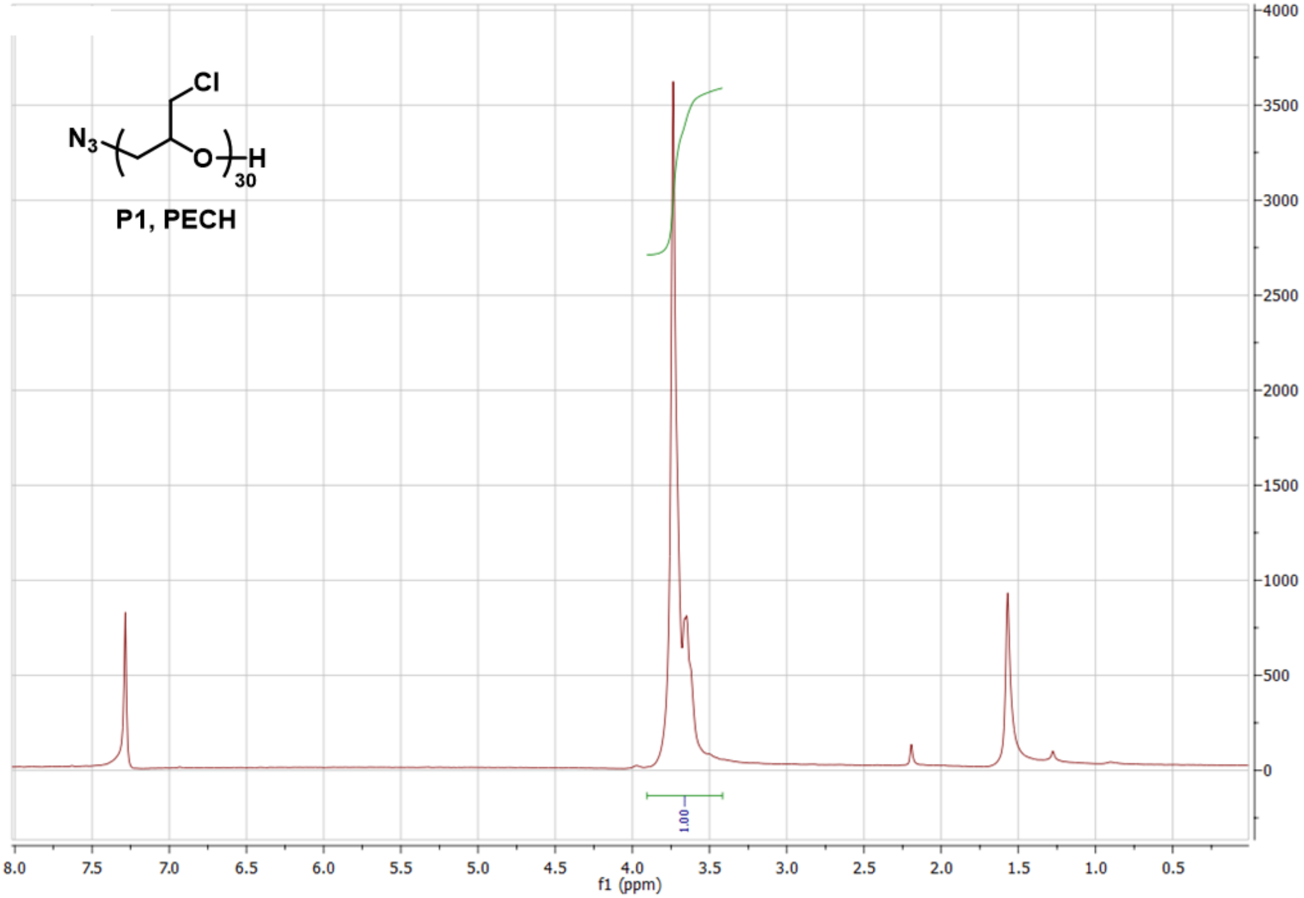

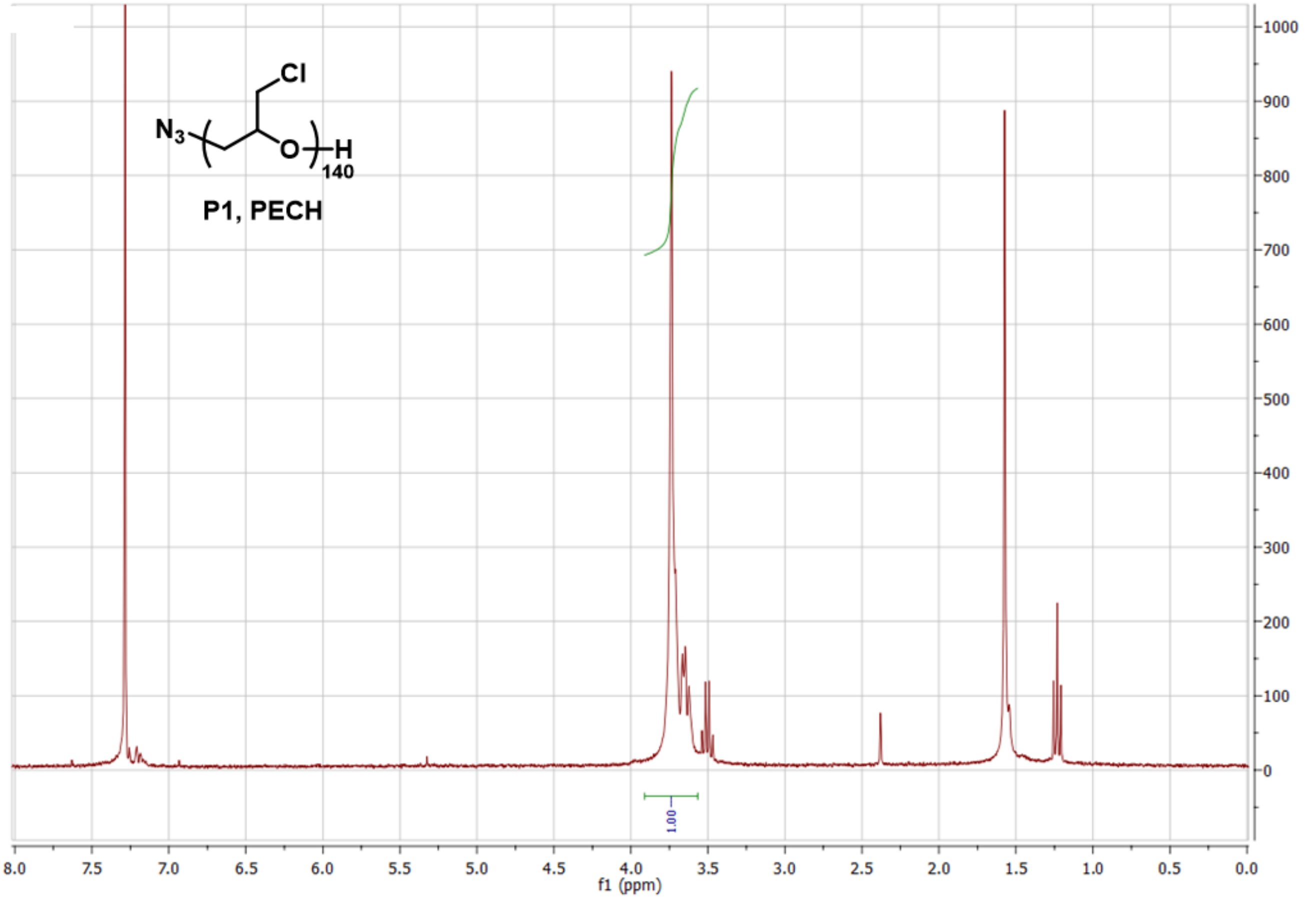
Cl
$N_3$
O
H
140
P1, PECH
1.00
1000
900
800
700
600
500
400
300
200
100
0
8.0
7.5
7.0
6.5
6.0
5.5
5.0
4.5
4.0
3.5
3.0
2.5
2.0
1.5
1.0
0.5
0.0
f1 (ppm)

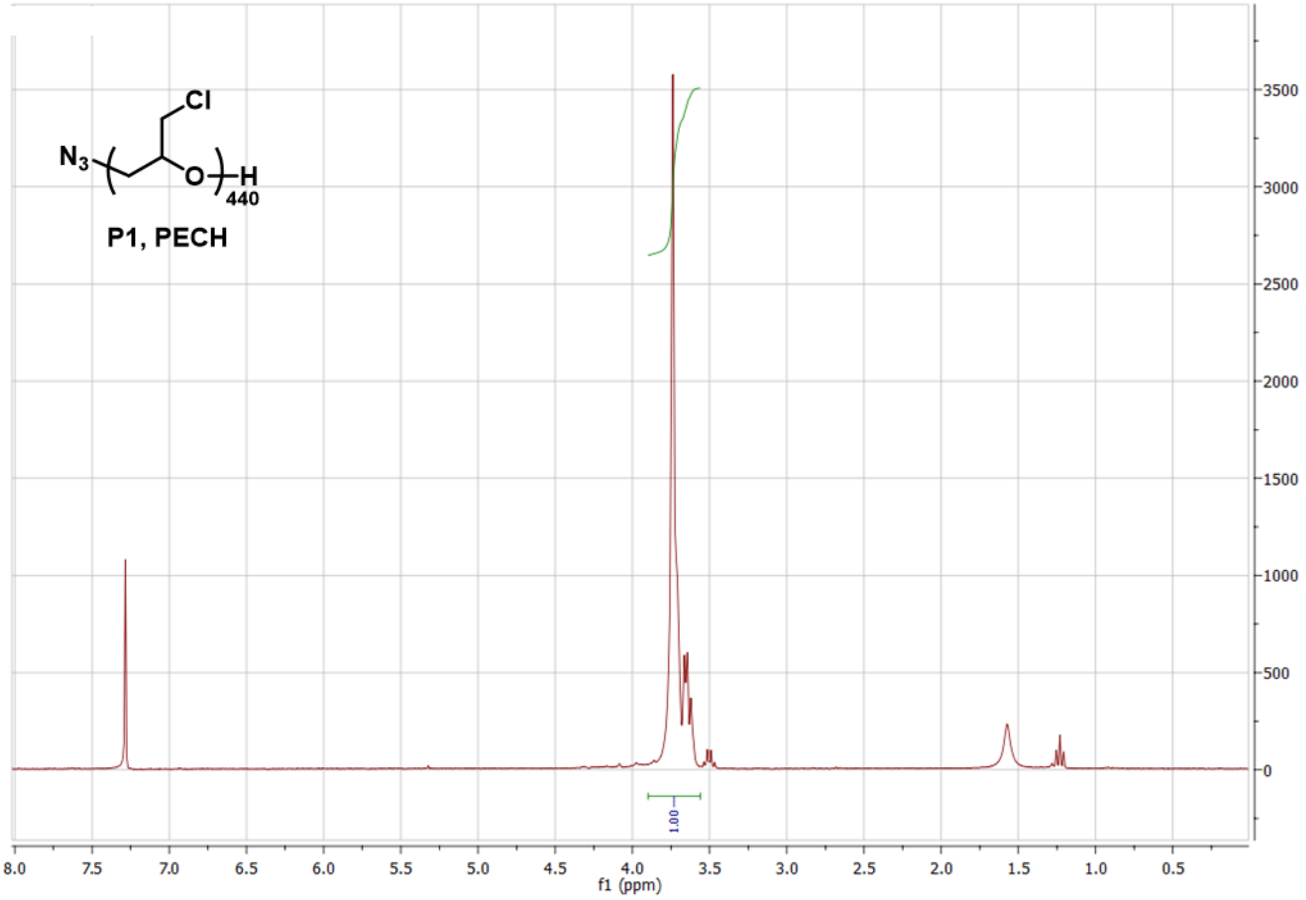

Cl
N3
O
H
440
P1, PECH
1.00
f1 (ppm)

**Figure S5**. $^{1}H$ NMR (300 MHz, $CDCl_3$) of cholestanone-terminated pECH polymers **P2-S/M/L**.

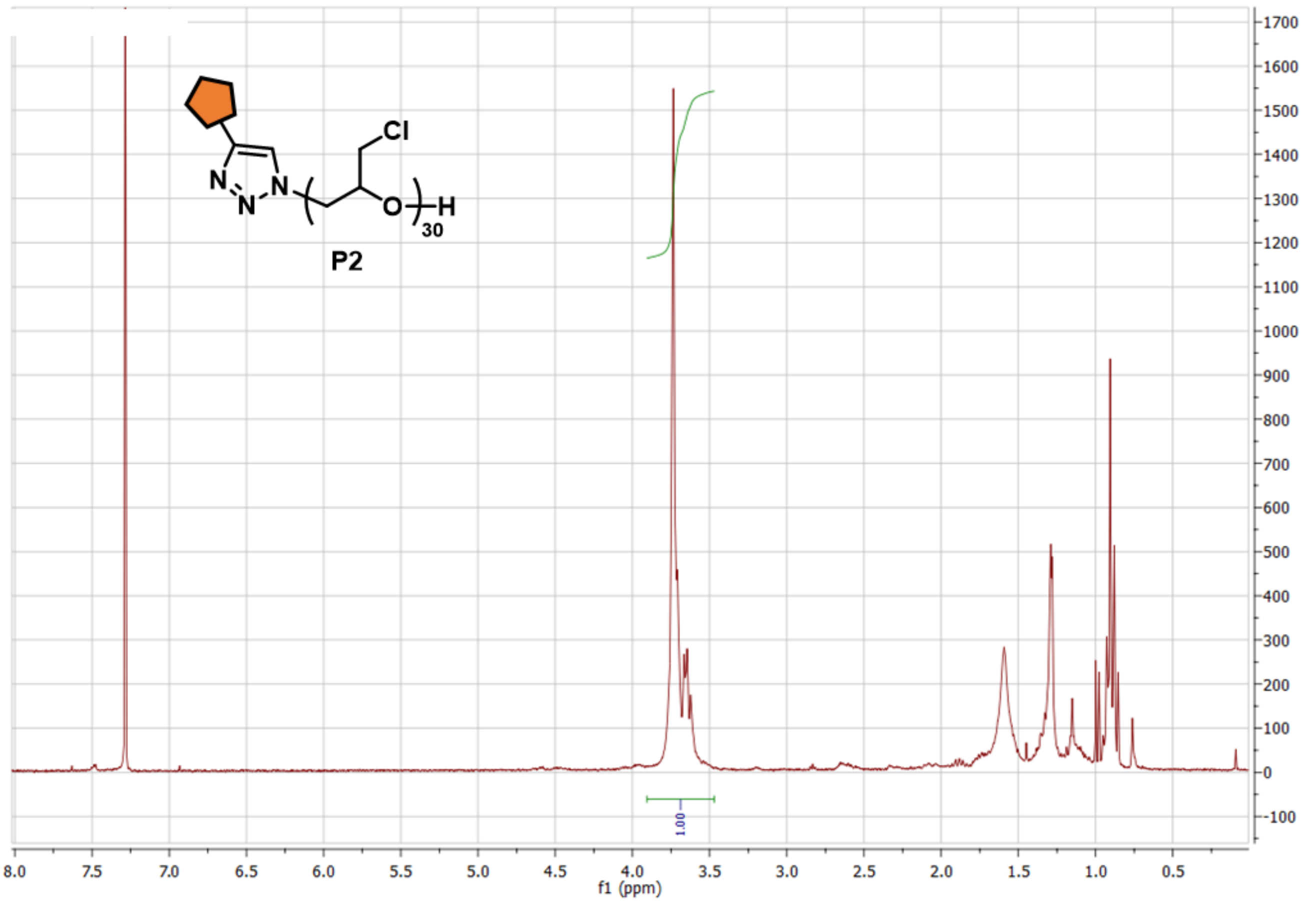

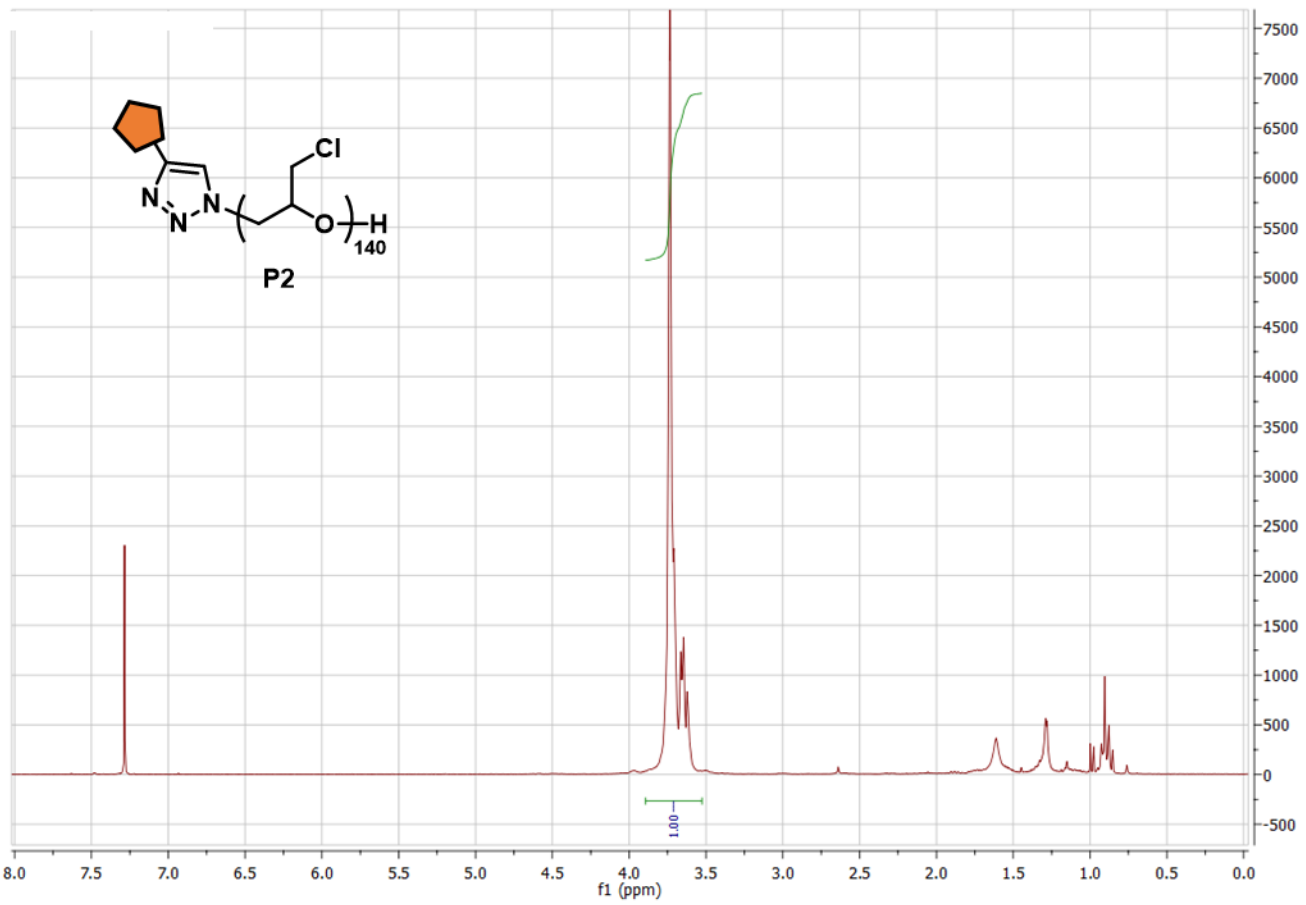

Cl
N
N
N
O
H
140
P2
1.00
8.0
7.5
7.0
6.5
6.0
5.5
5.0
4.5
4.0
3.5
3.0
2.5
2.0
1.5
1.0
0.5
0.0
f1 (ppm)
7500
7000
6500
6000
5500
5000
4500
4000
3500
3000
2500
2000
1500
1000
500
0
-500

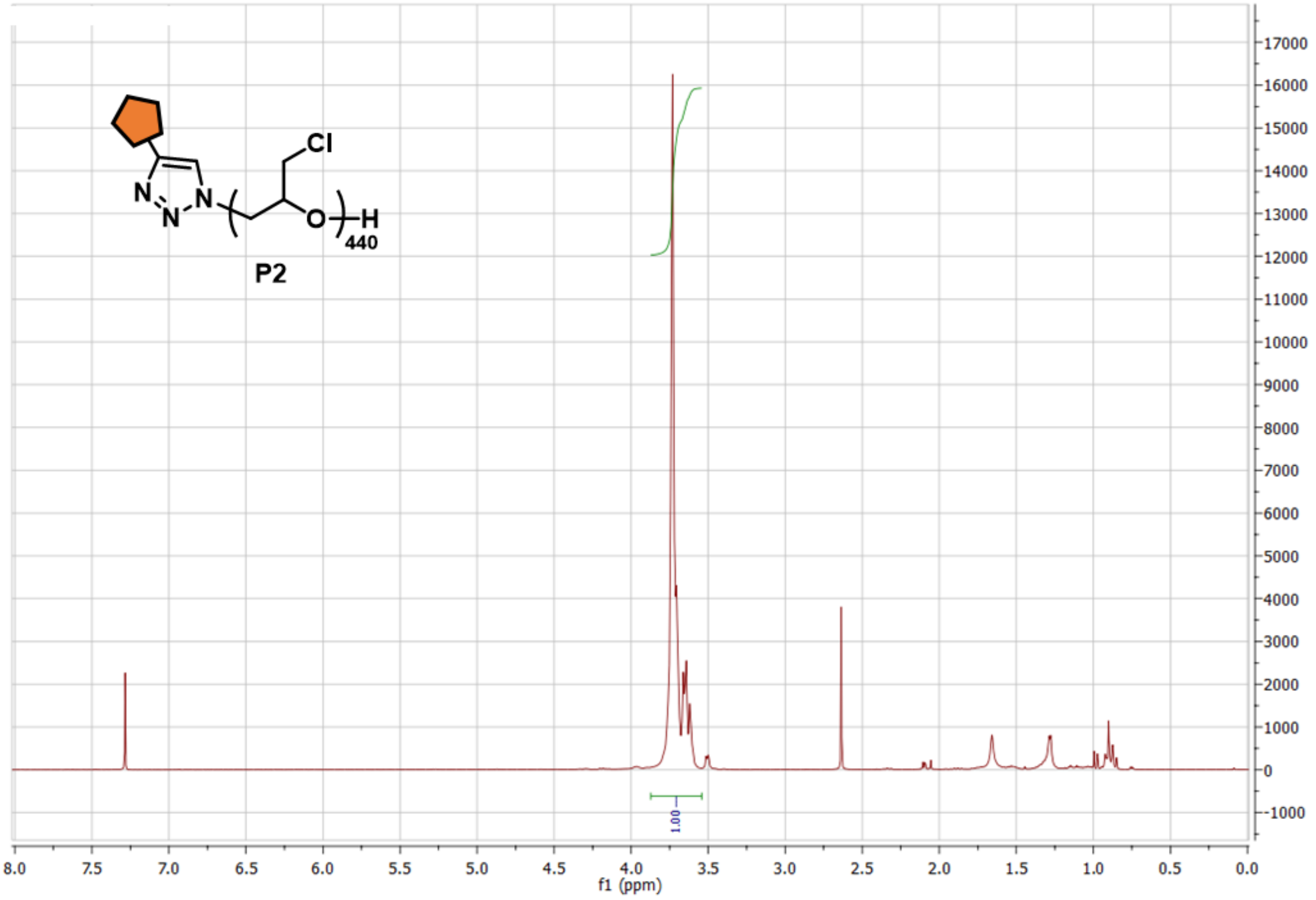

Cl
N
N
N
N
O
H
440
P2
17000
16000
15000
14000
13000
12000
11000
10000
9000
8000
7000
6000
5000
4000
3000
2000
1000
0
-1000
1.00
8.0
7.5
7.0
6.5
6.0
5.5
5.0
4.5
4.0
3.5
3.0
2.5
2.0
1.5
1.0
0.5
0.0
f1 (ppm)

**Figure S6**. [1]H NMR (300 MHz, $CDCl_3$) of cholestanone-terminated pGA polymers **P3-S/M/L**.

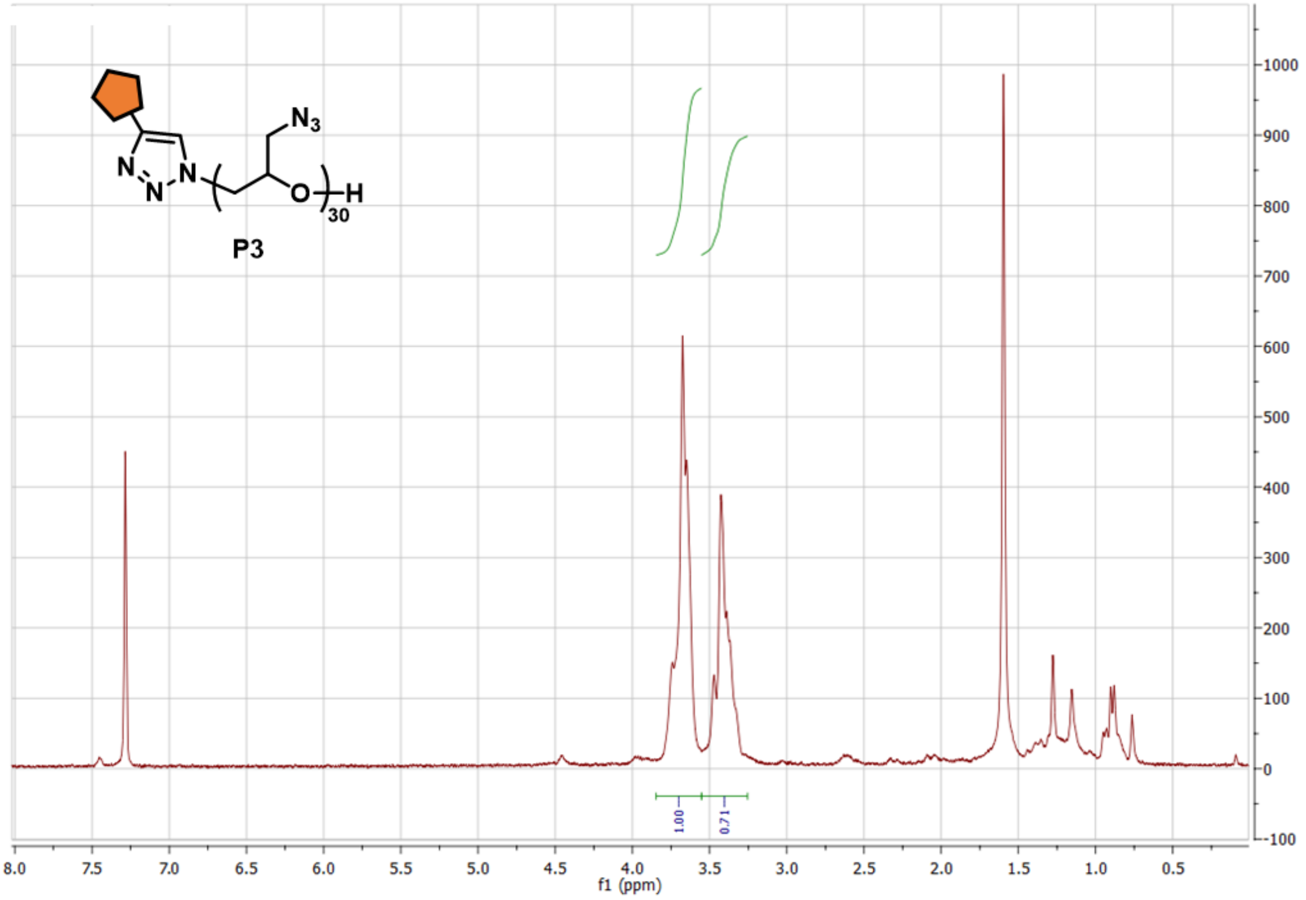

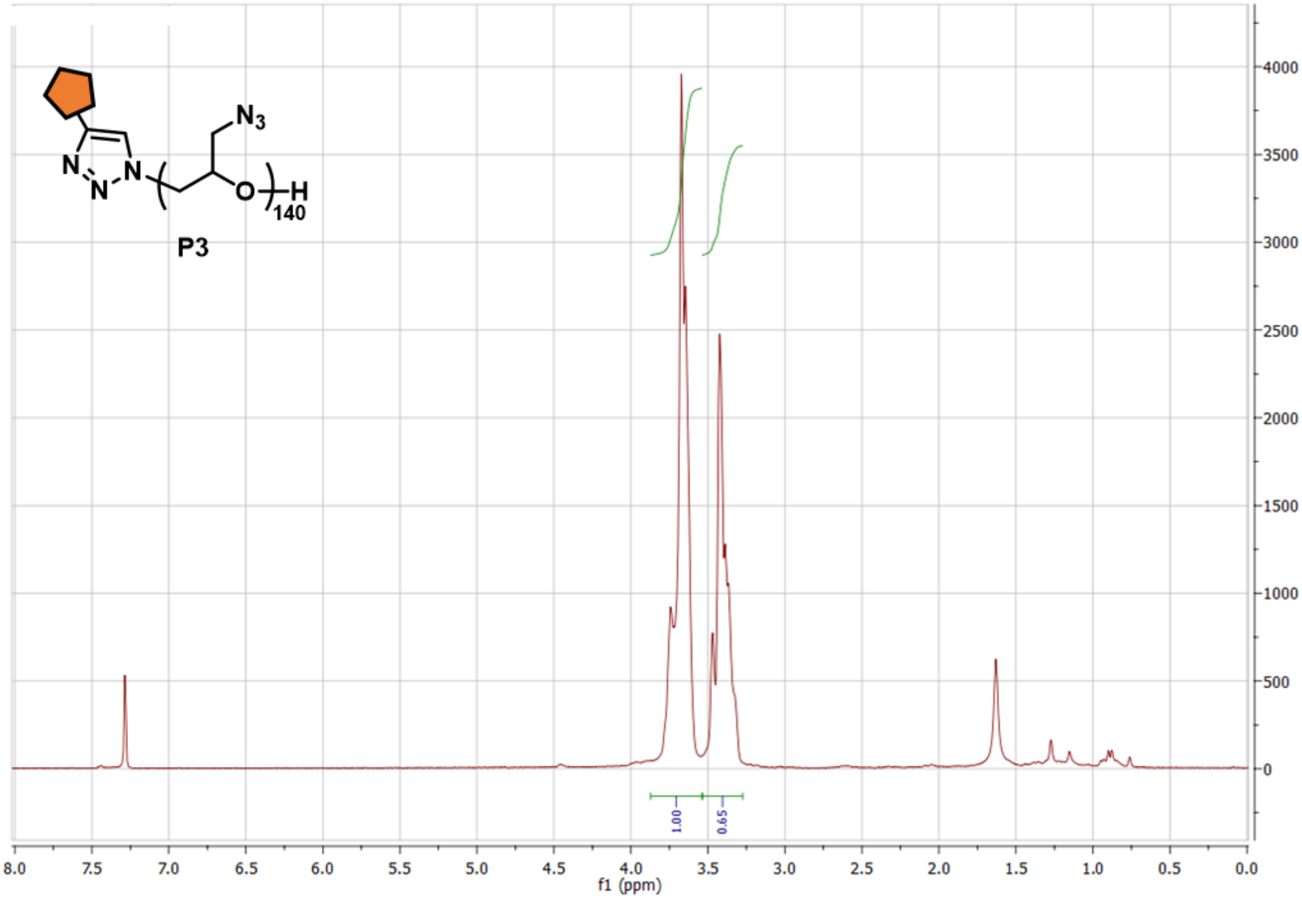
N3
N
N
N
140
H
O
P3
4000
3500
3000
2500
2000
1500
1000
500
0
1.00
0.65
8.0 7.5 7.0 6.5 6.0 5.5 5.0 4.5 4.0 3.5 3.0 2.5 2.0 1.5 1.0 0.5 0.0
f1 (ppm)

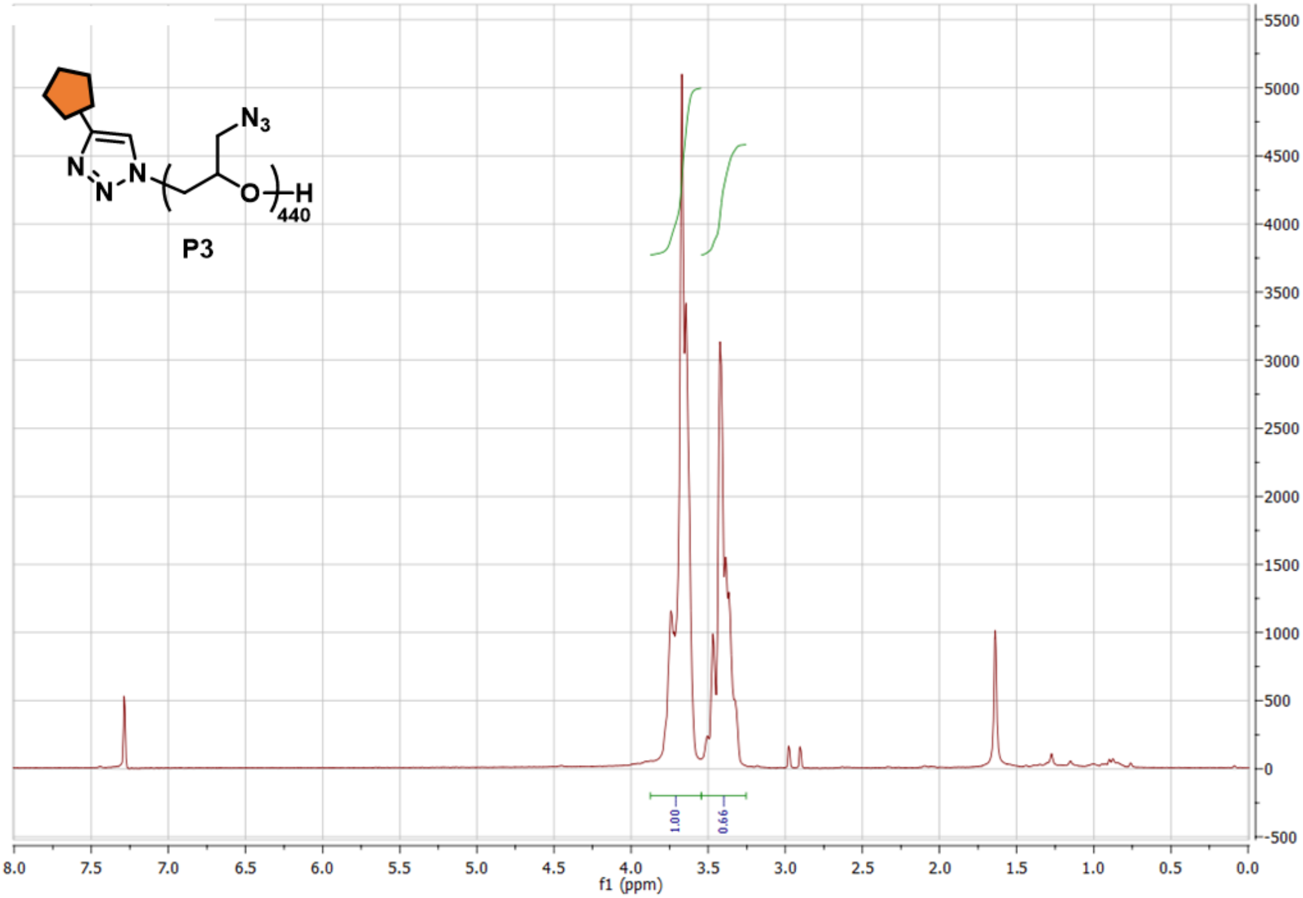

N3
440
P3
1.00
0.66
f1 (ppm)

**Figure S7**. $^{1}$H NMR (300 MHz, $D_2O$) of glycopolymers **GP-S/M/L**.

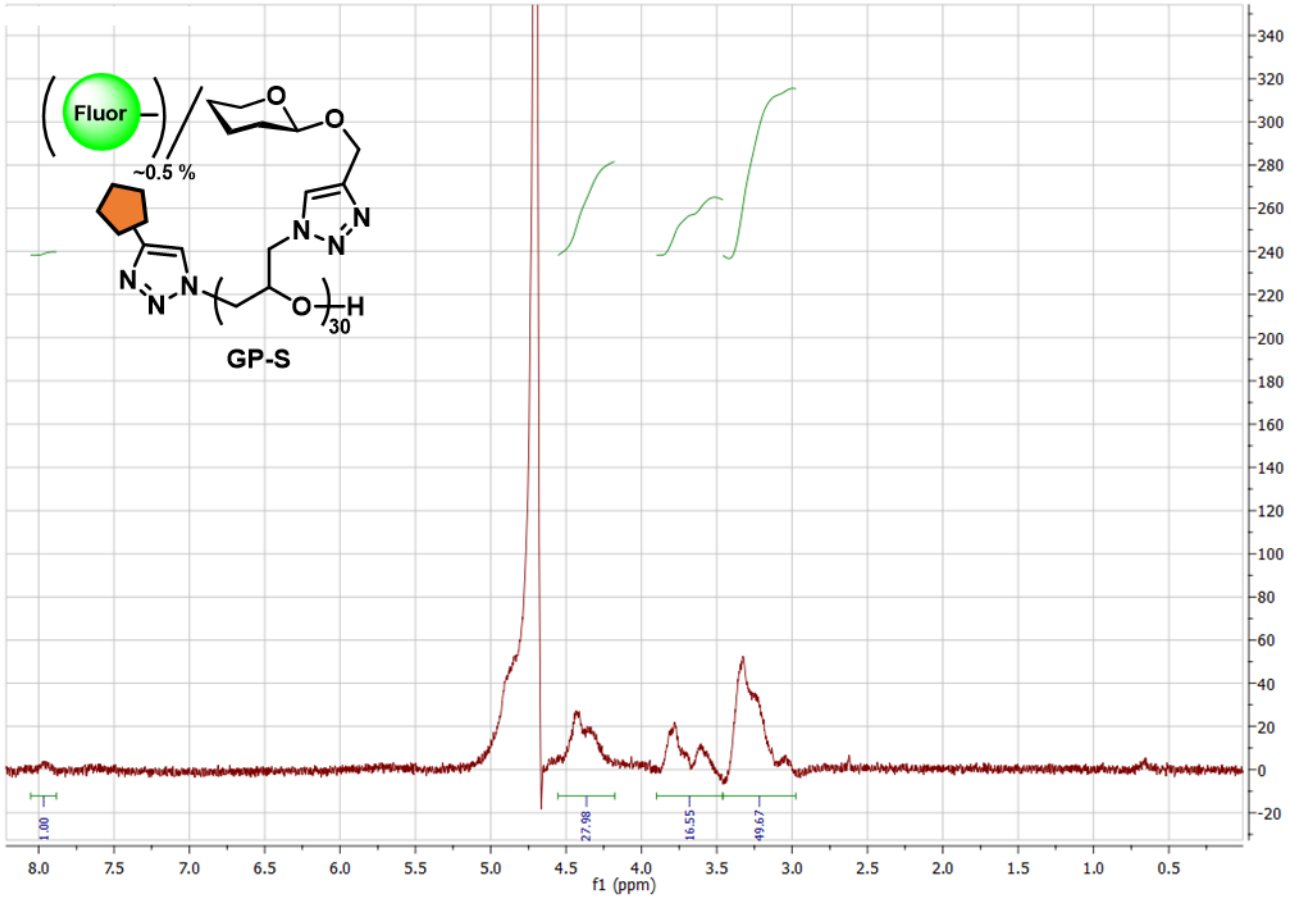

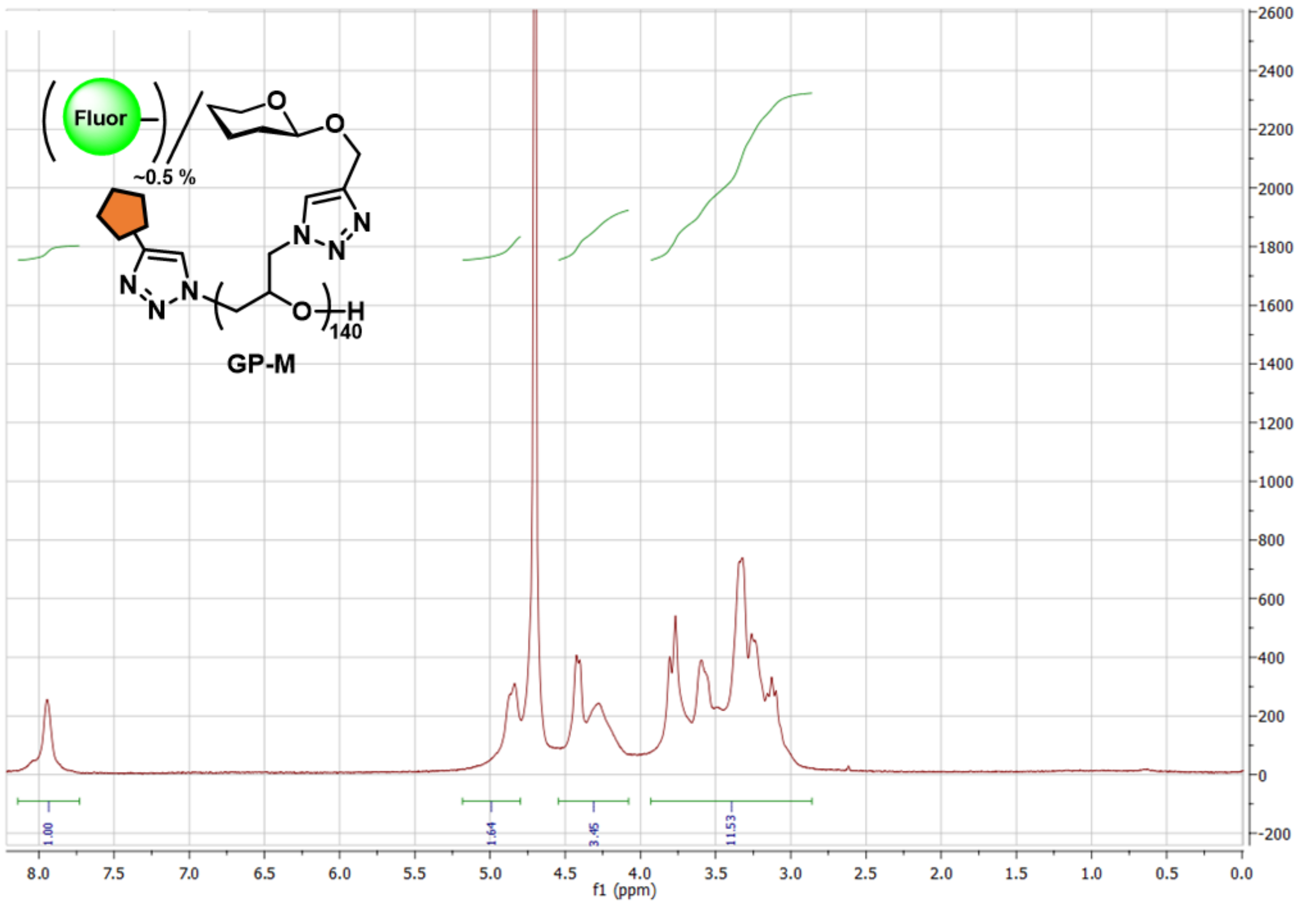
Fluor
~0.5 %
140
GP-M
1.00
1.64
3.45
11.53
f1 (ppm)

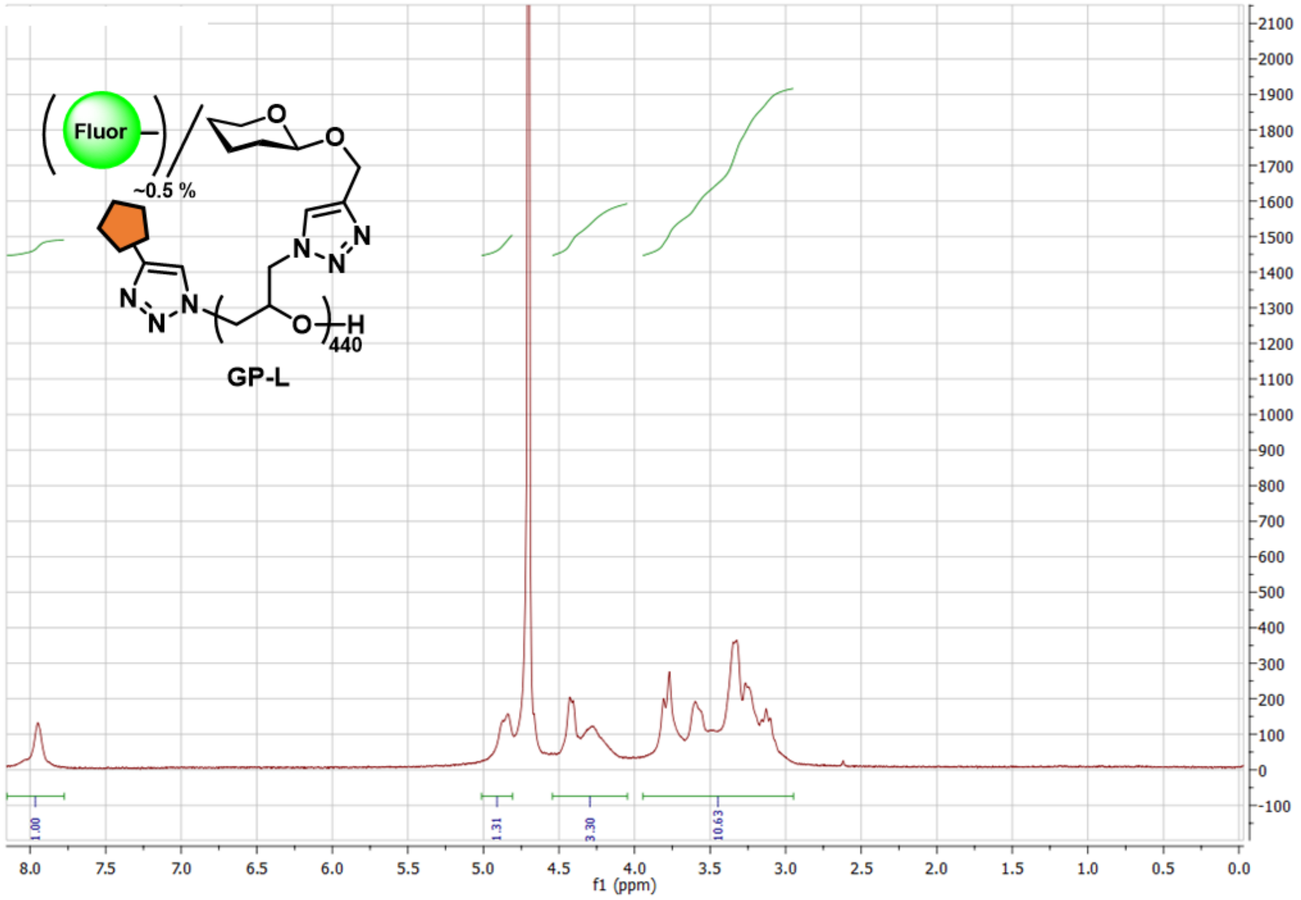
Fluor
~0.5 %
GP-L
440
1.00
1.31
3.30
10.63
f1 (ppm)

### Characterization of polymers P1, P2, P3, and GP-S/M/L by GPC and IR.

**Figure S8**. IR spectra of polymers **P1**, **P2**, **P3**, and **GP-S/M/L**.

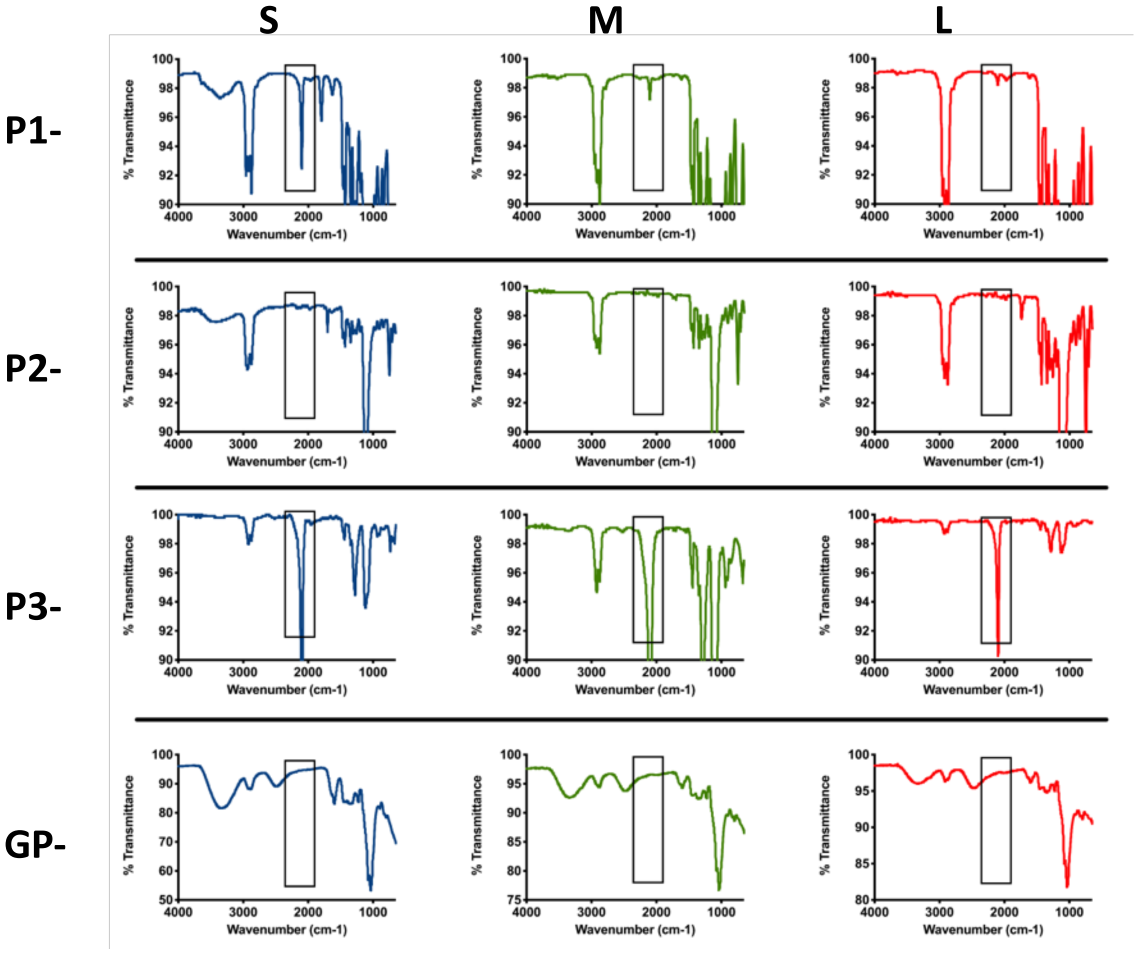

**Figure S9.** GPC traces for polymer intermediates (expanded data for **Fig 2A**).

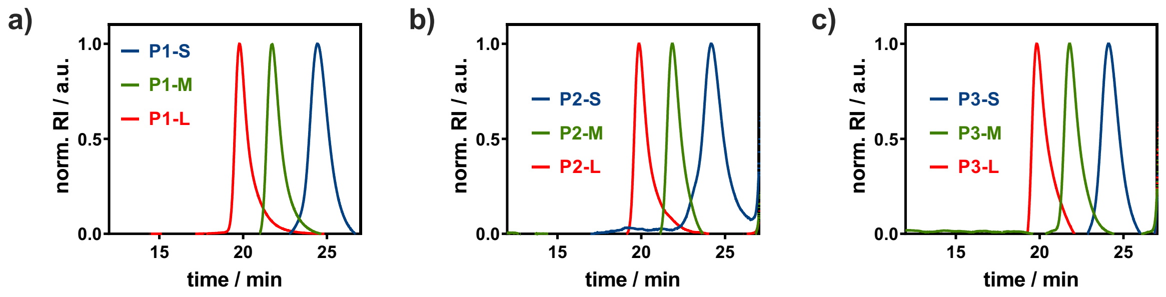

| | Polymerization conditions | | | | p(ECH) P1 | | P2 | | P3 | |
|---|---|---|---|---|---|---|---|---|---|---|
| length | [*i*-Bu3Al]/[NBu4N3] | [ECH] (n/L) | $Mn_{th}$ (kDa) | DP | Mn (kDa) | Đ | Mn (kDa) | Đ | Mn (kDa) | Đ |
| **S** | 1.5 | 2 | 3.0 | 30 | 3.1 | 1.32 | 3.5 | 1.3 | 5.8 | 1.23 |
| **M** | 2 | 2 | 10.0 | 140 | 12.7 | 1.19 | 13.7 | 1.27 | 19.9 | 1.11 |
| **L** | 3 | 2 | 45.0 | 440 | 40.5 | 1.23 | 45.3 | 1.21 | 51.1 | 1.22 |

**Table S10.** Expanded polymer characterization table for final Glycopolymers **GP-S/M/L.**

| Polymer (P) | DP | Mn (kDa) | Fl | Fl/P | Fl/DP | Glucose (%) |
|---|---|---|---|---|---|---|
| **GP-S** | 30 | 9.5 | AF488 | 0.3 | 1 | 100 |
| **GP-M** | 140 | 44.4 | AF488 | 0.86 | 0.61 | 100 |
| **GP-L** | 440 | 139.5 | AF488 | 2.7 | 0.61 | 100 |
| **GP-L/D** | 440 | 139.5 | AF488 | 2.4 | 0.55 | 100 |
| **GPL-L/A** | 440 | 139.5 | Cy3 | 2.7 | 0.61 | 100 |

Mn was calculated assuming 100% sidechain substitution, and using the theoretical DP attained from the parent pECH polymer **P1** (Mn = (Mw glycosylated side chain) x DP). Fluorophore labeling was quantified using the maximum absorbance values from UV/Vis. Glucose attachment was estimated to be quantitative because of the lack of an azide peak at 2100 $cm^{-1}$ in IR (**Fig S8**).

**RBC remodeling with glycopolymers GP-S/M/L.**

**Figure S11.** Relative levels of cell surface incorporation of glycopolymers **GP-S/M/L** at 7.5 µM.

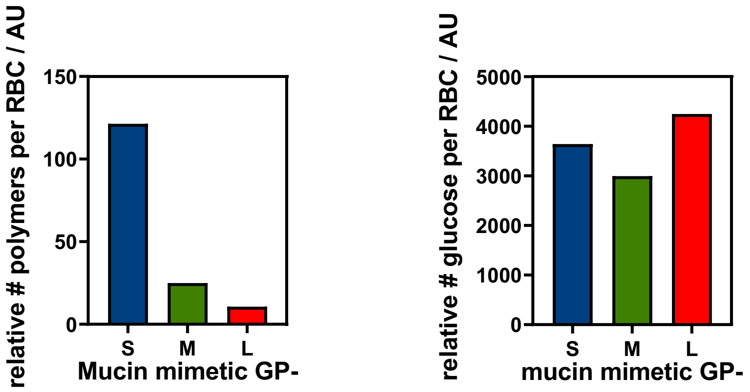

**Figure S12**. Relative incorporation of **GP-L** polymers vs equivalent polymer without cholestanone

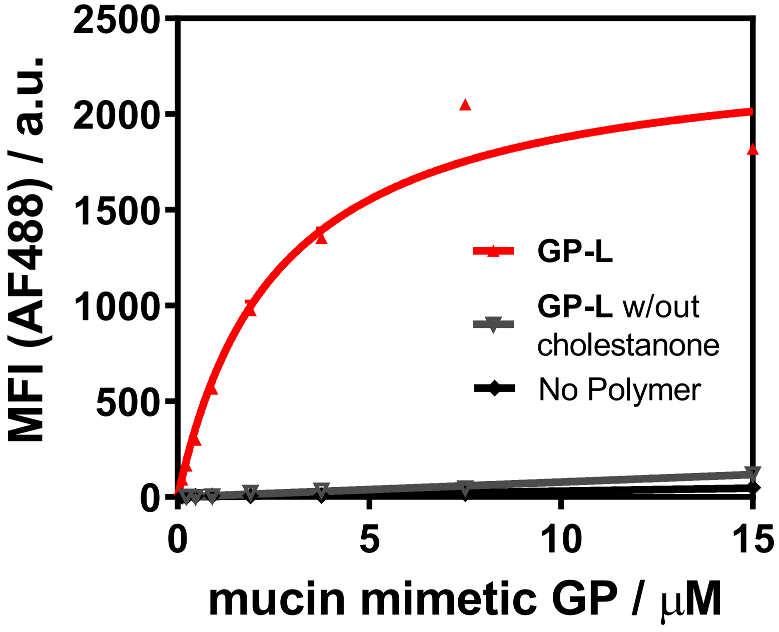

**Figure S13**. FSC and SSC of RBCs remodeled with glycopolymers **GP-S/M/L.**

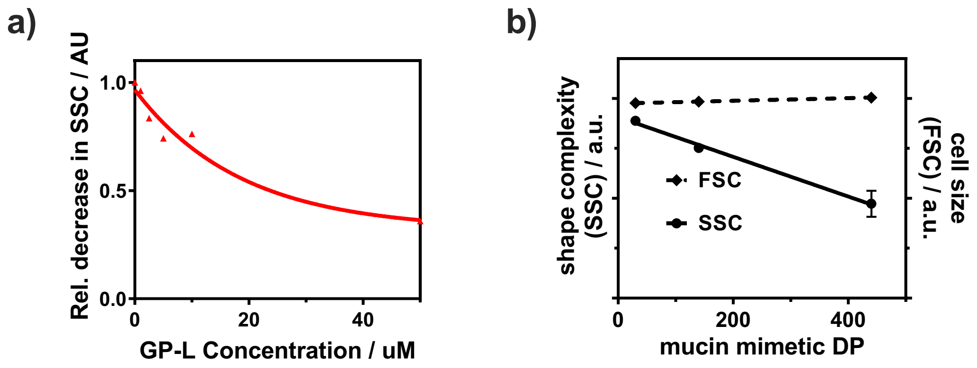

**Figure S14**. FRAP images of RBCs remodeled with **GP-S/M/L** (associated with **Fig 2F**).

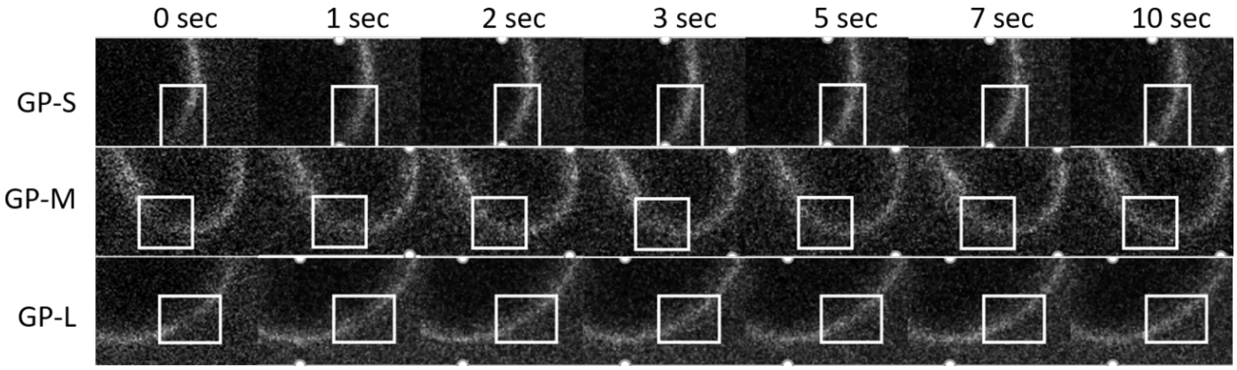

**Characterization of SNA binding to RBCs remodeled with glycopolymers GP-S/M/L.**

**Figure S15**. Binding of SNA to RBCs after pre-treatment with alkynyl cholestanone **S5.**

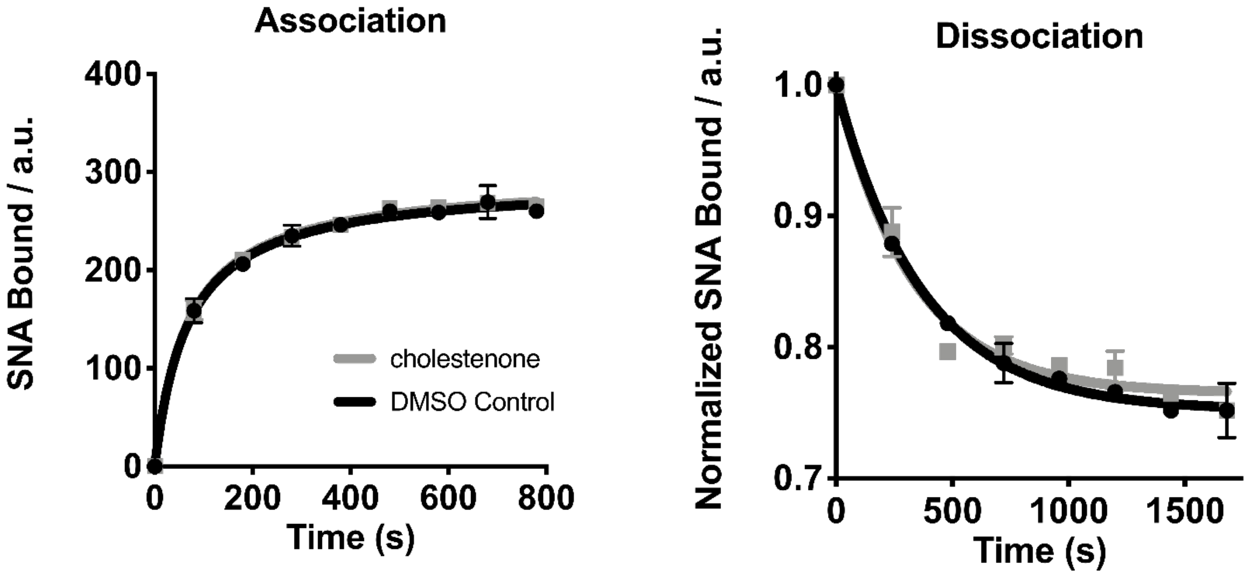

**Figure S16**. SNA binding to remodeled RBCs as a function of lectin concentration.

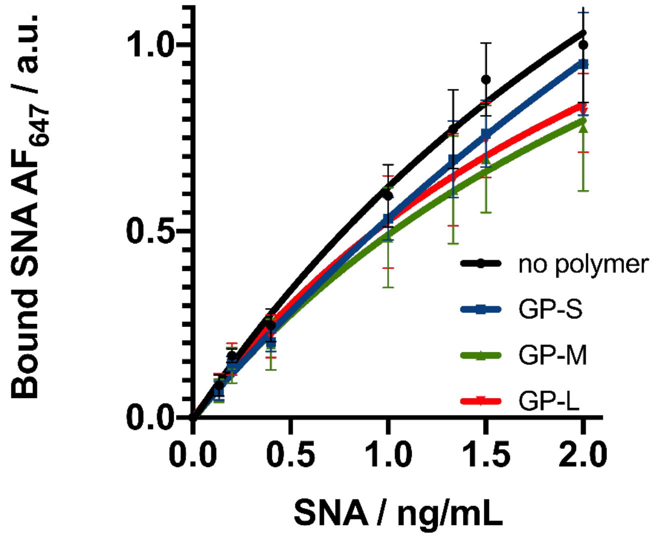

| SNA ng/mL | no polymer vs. GP-S | | no polymer vs. GP-M | | no polymer vs. GP-L | |
|---|---|---|---|---|---|---|
| | significance | p-value | significance | p-value | significance | p-value |
| 0.0 | ns | 0.9999 | ns | 0.9999 | ns | >0.9999 |
| 0.1 | ns | 0.9774 | ns | 0.9744 | ns | 0.9995 |
| 0.2 | ns | 0.9181 | ns | 0.8719 | ns | 0.992 |
| 0.4 | ns | 0.5649 | ns | 0.4702 | ns | 0.822 |
| 1.0 | ns | 0.3175 | * | 0.0207 | ns | 0.2222 |
| 1.3 | ns | 0.1364 | *** | 0.0003 | ** | 0.0037 |
| 1.5 | ** | 0.0018 | **** | <0.0001 | *** | 0.0004 |
| 2.0 | ns | 0.4641 | **** | <0.0001 | **** | <0.0001 |

**Figure S17**. Mobility of SNA bound to RBC membrane (FRAP).

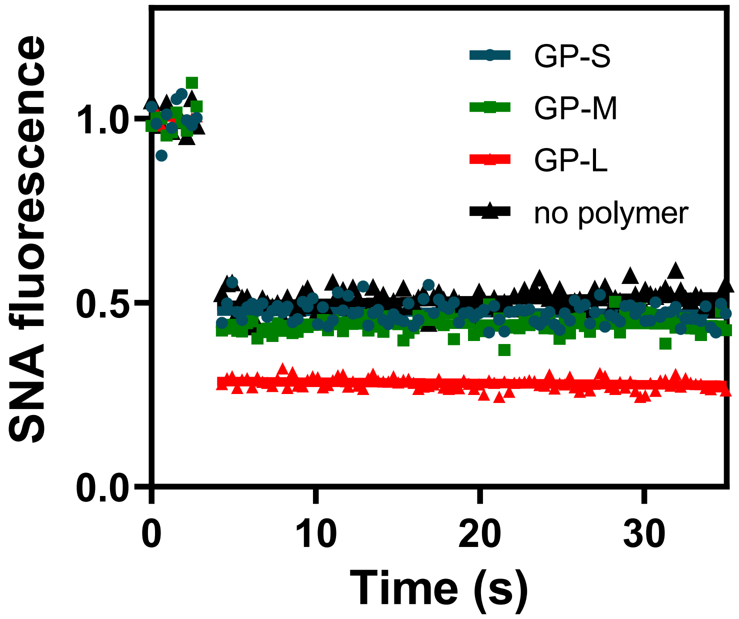

**H1N1 binding to RBCs remodeled with glycopolymers GP-S/M/L via enzymatic 4MU-NANA assay.**

**Figure S18**. 4MU-NANA fluorescence turn on with increasing viral titer.

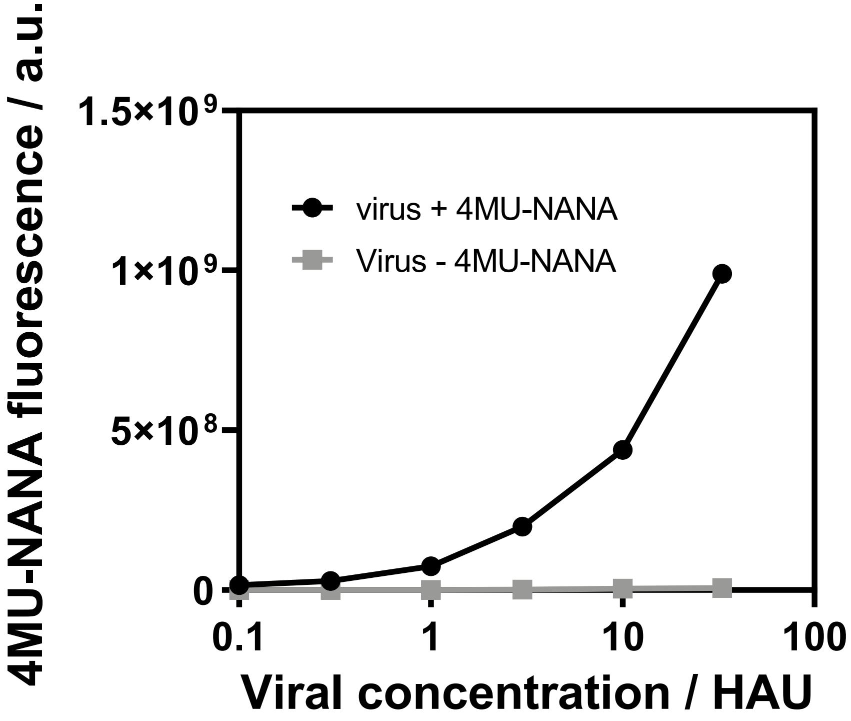

**Figure S19**. Viral titer dependance on binding to RBCs.

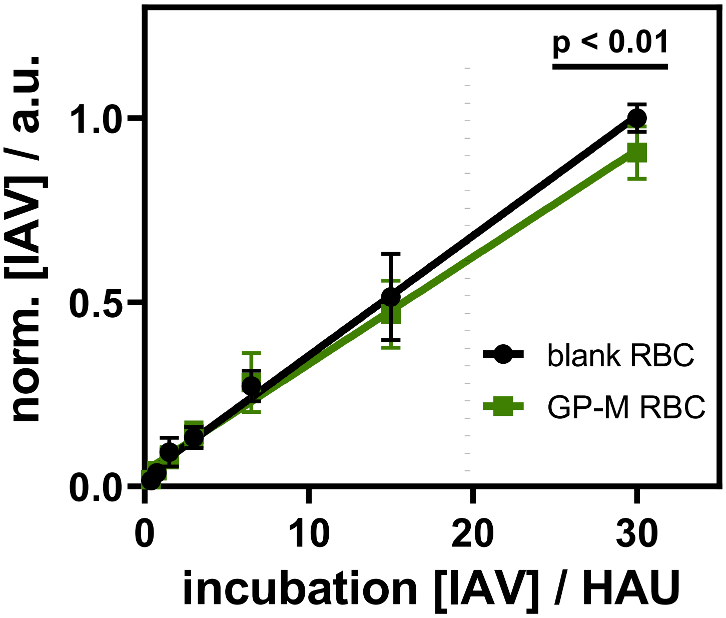
